# The CorC proteins MgpA (YoaE) and CorC protect from excess-cation stress and are required for egg white tolerance and virulence in *Salmonella*

**DOI:** 10.1101/2025.03.18.643926

**Authors:** Yumi Iwadate, James M. Slauch

## Abstract

Cation homeostasis is a vital function. In *Salmonella*, growth in very low Mg^2+^ induces expression of high-affinity Mg²⁺ transporters and synthesis of polyamines, organic cations that substitute for Mg²⁺. Once Mg²⁺ levels are re-established, the polyamines must be excreted by PaeA. Otherwise, cells lose viability due to a condition we term excess-cation stress. We sought additional tolerance mechanisms for this stress. We show that CorC and MgpA (YoaE) are essential for survival in stationary phase after Mg^2+^ starvation. Deletion of *corC* causes a loss of viability additive with the *paeA* phenotype. Deletion of *mgpA* causes a synthetic defect in the *corC* background. This lethality is suppressed by loss of the inducible Mg^2+^ transporters, suggesting that the *corC mgpA* mutant is sensitive to changes in intracellular Mg^2+^. CorC and MgpA function independently of PaeA. A *paeA* mutant is sensitive to externally added polyamine in stationary phase; loss of CorC and MgpA suppressed this sensitivity. Conversely, the *corC mgpA* mutant, but not the *paeA* mutant, exhibited sensitivity to high Mg^2+^ and egg white. The *corC mgpA* mutant is also attenuated in a mouse model. The *corC* and *mgpA* genes are induced in response to increased Mg^2+^ concentrations. Thus, CorC and MgpA play some interrelated role in cation homeostasis. It is unlikely that these phenotypes are due to absolute levels of cations. Rather, the cell maintains relative concentrations of various cations that likely compete for binding to anionic components. Imbalance of these cations affects some essential function(s), leading to a loss of viability.

**IMPORTANCE:** Mg²⁺ and other cations are critical for counteracting anionic compounds in the cell including RNA, DNA, and nucleotides. Both excessively low and high cation levels are toxic. To maintain proper intracellular concentrations, cells must regulate Mg²⁺ importers and exporters, or modulate the levels of other cations or anions that affect free Mg²⁺ levels. In *Salmonella*, no mutants sensitive to high Mg²⁺ levels have been identified. Here, we demonstrate that the largely uncharacterized proteins MgpA and CorC are induced under high Mg²⁺ conditions and are essential for tolerance to high Mg²⁺ levels. These genes are also essential for survival during endogenous excess-cation stress triggered by the transition to stationary phase after Mg²⁺ starvation, as well as for virulence, highlighting the broader role of cation homeostasis.

## INTRODUCTION

Cations, particularly magnesium (Mg^2+^), are required to stabilize RNA and DNA (1–3), counteract the negative charges on ATP (4), and neutralize the negative charges in outer membrane lipopolysaccharide (LPS) (5). Mg^2+^ is also critical for ribosome assembly, function, and stability (4, 6). Polyamines are organic cations important in all domains of life, but their physiological role is not well understood. Our recent data support a new paradigm for the overall role of polyamines in cell physiology. Our primary hypothesis is that polyamine and Mg^2+^ concentrations are coordinately controlled in the cell and that polyamines act as simple cations that can substitute for Mg^2+^ in critical systems under low-Mg^2+^ stress.

*Salmonella* Typhimurium is a major food-borne pathogen that replicates within macrophages, and adaptation to the low Mg^2+^ environment of the macrophage phagosome is key to *Salmonella* virulence. Counterintuitively, shifting *Salmonella* cells to Mg^2+^ deplete conditions induces an ultimate overload of cations that can be lethal upon reaching stationary phase. Upon Mg^2+^ starvation, polyamine synthesis is induced (7, 8), as is production of high affinity Mg^2+^ transporters MgtA and MgtB (9–13). Either polyamine synthesis or Mg^2+^ transport is required to maintain viability under these low Mg^2+^ conditions. Once Mg^2+^ levels are re-established, the excess polyamines must be excreted by the inner membrane polyamine exporter PaeA (7, 14). Mutants lacking PaeA lose viability when cells reach stationary phase. The lethality of the *paeA* mutant in stationary phase is suppressed by blocking Mg^2+^ transport, indicating that it is the total concentration of Mg^2+^ and polyamines that is detrimental, a phenomenon that we term “excess-cation stress.” Importantly, these results are recapitulated during infection. Polyamine synthesis mutants are attenuated in a mouse model of systemic infection, as are strains lacking the MgtB Mg^2+^ transporter. Combined, these mutations confer a synthetic phenotype in animals, confirming that Mg^2+^ and polyamines are required for the same overall processes. These data support our hypothesis that the cell coordinately controls polyamine and Mg^2+^ concentrations to maintain overall cation homeostasis.

We sought additional proteins involved in mitigating excess-cation stress in *Salmonella*. We focused on the domain structure of the PaeA protein, particularly its C-terminal CorC domain. The PaeA protein contains an N-terminal transmembrane CNNM domain (pfam01595), and a cytoplasmic portion containing a CBS pair domain (pfam00571), followed by the C-terminal CorC_HlyC domain (pfam03471). The *Salmonella* genome encodes five additional proteins with CorC domains: CorB, CorC, MgpA (YoaE), YegH, and CvrA (Fig 1). CorB resembles PaeA with a CNNM transmembrane domain, CBS pair, and C-terminal CorC domain. The eponymous CorC lacks a transmembrane domain and has only a CBS pair and C-terminal CorC domain. MgpA and YegH each possess an N-terminal transmembrane TerC domain (pfam03741) followed by a CBS pair and C-terminal CorC domain. The CvrA protein contains an N-terminal transmembrane Na/H^+^ exchanger domain (pfam00999), and a cytoplasmic RCK domain (pfam02080), followed by a CorC domain.

**Fig. 1.**
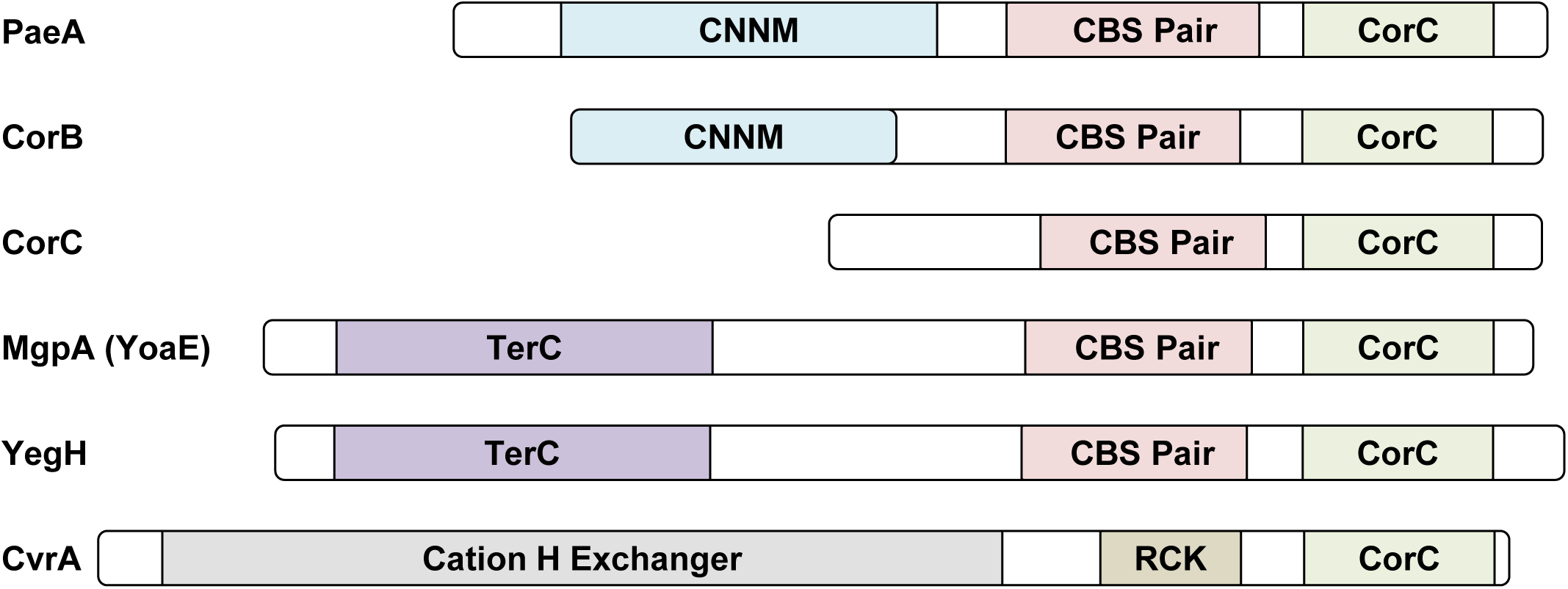
Domain architecture of CorC domain-containing proteins in *Salmonella*. Conserved domains were predicted using InterPro (https://www.ebi.ac.uk/interpro/). Each domain is represented by a different color, while regions with no domain assignment are shown in white.

Although widespread, the biological and biochemical function of the CorC domain remains unknown. We postulate that the CorC domain or the CBS pair/CorC domains function as a sensor of cytoplasmic cations, such as Mg^2+^. For example, our model suggests that PaeA exports polyamines in response to the Mg^2+^ concentration (7). Moreover, all characterized CorC domain-containing proteins appear to be involved in cation transport. PaeA in *Salmonella* and *E. coli* is responsible for the efflux of cadaverine and putrescine (14). CorC and CorB were originally identified as conferring resistance to cobalt in the medium and were further implicated in Mg^2+^ efflux (15). CorB homologs in the thermophilic archaeon *Methanoculleus thermophilus* and the thermophilic Gram-negative bacterium *Tepidiphilus thermophilus* are well-characterized Mg^2+^ efflux transporters (16, 17). In *Staphylococcus aureus*, the CorB homolog, MpfA, is essential for tolerance to high Mg^2+^ levels (18). The CvrA protein in *Vibrio cholerae* facilitates K^+^/H^+^ exchange (19). Collectively, these reports suggest that CorC domain-containing proteins are crucial for maintaining proper cation levels and relative composition.

Given these findings, we examined whether any of the CorC domain-containing proteins in *Salmonella* play a critical role in reducing excess-cation stress. Our results show that two of these proteins, CorC and MgpA (YoaE), are required for full survival during stationary phase after Mg^2+^ starvation, for full virulence in a mouse model, and for tolerance to high Mg^2+^ stress. (We have renamed *yoaE* to *mgpA* for <u>Mg</u>^2+^ <u>p</u>rotection protein <u>A</u>). Thus, both CorC and MgpA play roles in mitigating excess-cation stress, whether it arises endogenously or exogenously. Moreover, CorC and MgpA are required for egg white tolerance, suggesting that exposure to egg white imposes cation imbalance in *Salmonella*. The constitutive Mg^2+^ transporter CorA, previously believed to genetically interact with CorC (15), is dispensable for CorC’s role in survival during stationary phase after Mg^2+^ starvation, showing that CorC functions independently of CorA. Our findings reveal that multilayered systems involving CorC domain-containing proteins maintain proper cation levels in response to environmental changes, including during host infection.

## RESULTS

### CorC is required for full survival under excess-cation stress in stationary phase

Our previous studies revealed that, upon Mg^2+^ starvation, *Salmonella* induces both polyamine synthesis and high affinity Mg^2+^ transport. Once Mg^2+^ levels are restored, PaeA is responsible for the efflux of the polyamines cadaverine and putrescine. In the absence of PaeA, cells start to die upon entry into stationary phase (7, 14). We seek to further understand this excess-cation stress in stationary phase.

The C-terminus of PaeA contains a CorC domain. To better understand the CorC domain, we initially focused on the eponymous CorC protein and investigated its role in response to induction of excess-cation stress. The wild-type and Δ*corC* strains were grown in N-minimal medium (pH 7.4) with or without Mg^2+^ at 37℃, and viability and OD_600_ were monitored for 48 hours. The Δ*corC* strain displayed a decline in viability upon entry into stationary phase after Mg^2+^ starvation (Fig 2A). The OD_600_ was unaffected in the Δ*corC* strain compared to the wild type grown in no Mg^2+^, showing that the cells were not lysing (Fig S1A). The *corC* gene is transcribed in an operon with upstream genes *ybeZ* and *ybeY* and downstream gene, *lnt* (20–22). To confirm that the phenotype results from loss of CorC, we introduced low-copy number plasmids pWKS30 (vector) or pWKS30-*corC* into the Δ*corC* strain and examined viability after Mg^2+^ starvation. As shown in Figure S2A, the phenotype of the Δ*corC* strain was fully complemented by pWKS30-*corC*. These results suggest that CorC has an important role in maintaining viability in stationary phase after Mg^2+^ starvation, similar to PaeA.

**Fig. 2.**
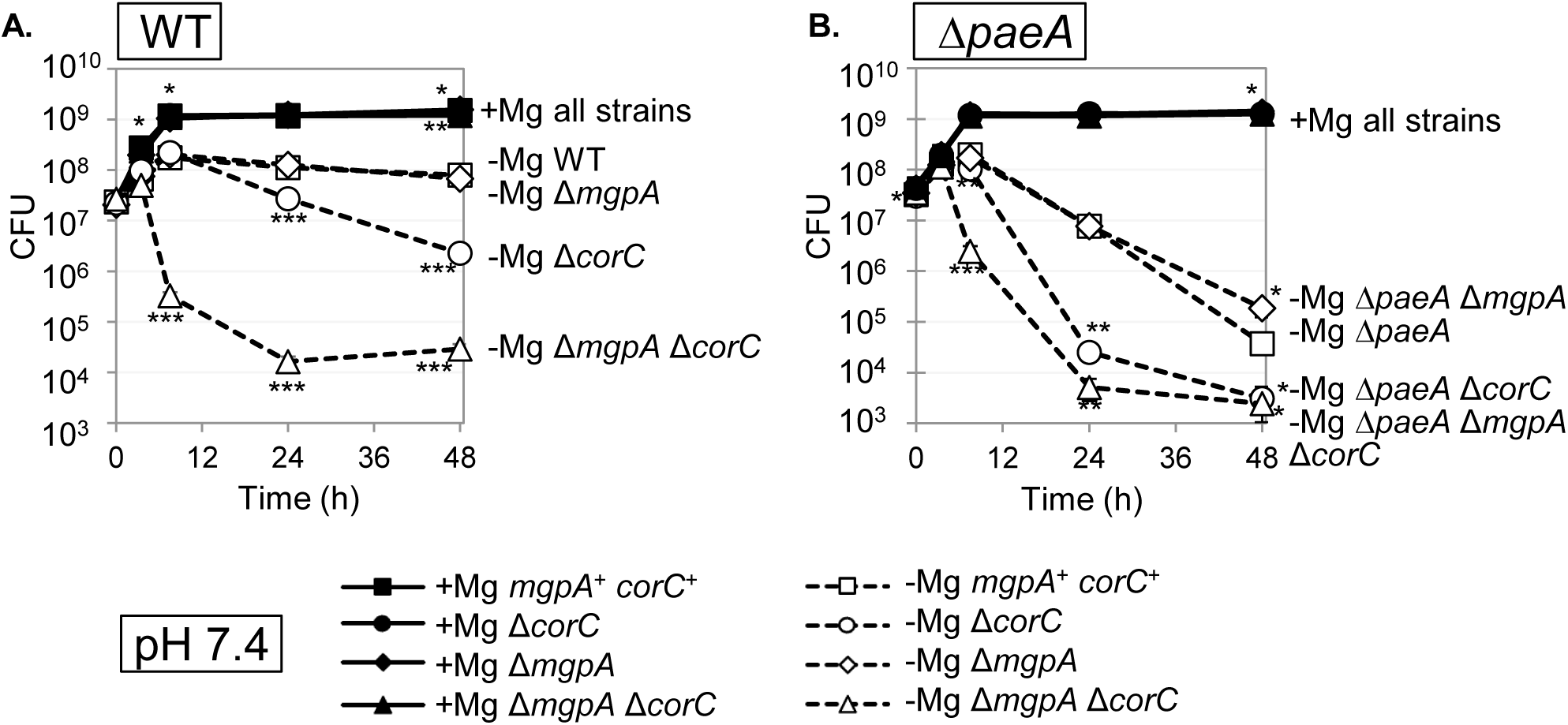
The Δ*corC* strain loses viability in stationary phase after Mg^2+^ starvation, the deletion of *mgpA* exacerbates the effect. The indicated strains were grown overnight in N-minimal medium pH 7.4 with 10 mM MgCl_2_, washed, diluted into N-minimal medium pH 7.4 with or without 10 mM MgCl_2_ (t=0 h), and incubated at 37℃. CFUs were determined at the indicated time points in wild type (A) and ΔpaeA (B) background. Values are mean ± SD, n = 6. Unpaired t test (*p* < 0.05*, 0.005**, 0.0005***) vs *mgpA*^+^ *corC*^+^ parent strain at the same time point and at the same Mg²⁺ concentration. Strains used: 14028, JS2692, JS2693, JS2694, JS2695, JS2696, JS2697, and JS2698.

### Loss of CorC and MgpA confers synthetic phenotypes

CorC is predicted to be a cytoplasmic protein with no transmembrane domain, suggesting that CorC is not a transporter per se. To investigate the mechanisms underlying the role of CorC in maintaining survival in stationary phase after Mg^2+^ starvation, we asked if there was any genetic interaction between CorC and other CorC domain-containing proteins. As we reported previously (7), a Δ*paeA* strain showed a loss of viability in stationary phase after Mg^2+^ starvation (Fig. 2B). The deletion of the *corC* gene in the Δ*paeA* background further decreased viability in stationary phase, with the results suggesting an apparent additive effect in the double mutant (compare Fig. 2A and B). This suggests that *paeA* and *corC* are independently involved in the same phenomenon, in a broad sense.

Deletion of *mgpA* had no apparent effect in stationary phase after Mg^2+^ starvation. However, deletion of *mgpA* in the *corC* background notably intensified the loss of viability upon transition to stationary phase after Mg^2+^ starvation (Fig. 2A). This suggests a synthetic relationship between the *mgpA* and *corC* genes. In contrast to *mgpA*, deletion of the remaining CorC domain-containing proteins, *corB, yegH, or cvrA*, conferred no apparent phenotype in either the wild-type or Δ*corC* backgrounds (Fig. S3), indicating that they are not involved in this CorC-related phenomena under these conditions. Note that YegH has the same general domain structure as MgpA (Fig. 1).

Introducing the pWKS30-*corC* plasmid into the Δ*mgpA* Δ*corC* strain restored viability in the stationary phase, while a pWKS30-*mgpA* plasmid restored the phenotype to that of the Δ*corC* strain (Fig. S2B). StyR-291, a sRNA encoded downstream of *mgpA* and antisense to the 3’-terminus of the *pdeD* gene (23), does not affect the phenotype; a Δ*mgpA* Δ*corC* strain harboring the pWKS30-*mgpA-* StyR-291 plasmid showed the same level of suppression of the strain harboring pWKS30-*mgpA* (Fig. S2B). Deleting *mgpA* had no phenotypic effect in the Δ*paeA* strain, showing that there is no genetic interaction between *mgpA* and *paeA* (Fig. 2B). The Δ*mgpA* mutation did exacerbate the phenotype seen in the Δ*corC* Δ*paeA* background, as expected (Fig. 2B).

### CorC and MgpA are required for survival when Mg^2+^ is imported

The lethality conferred by loss of PaeA after Mg^2+^ starvation can be suppressed by loss of MgtA and MgtB Mg^2+^ transporters or by blocking polyamine synthesis (Δsynth) (7). To gain further insight into the phenotypes of *mgpA* and *corC* mutants after Mg^2+^ starvation, we performed similar analyses. Loss of MgtA and MgtB completely suppressed the loss of viability conferred by the deletion of *corC* or of both *mgpA* and *corC* in stationary phase (Fig. 3A and B), mirroring the effect seen in the Δ*paeA* mutant (7). This similarity implies that the transport of Mg^2+^ via inducible Mg^2+^ transporters serves as a primary factor contributing to the loss of viability in these various mutants, consistent with the concept that Mg^2+^ starvation induces Mg^2+^ uptake leading to excess-cation stress. In contrast, the deletion of polyamine synthesis genes only partially suppressed the phenotypes in the Δ*corC* and the Δ*mgpA* Δ*corC* mutants (Fig. 3C), differing from the complete suppression observed in Δ*paeA* strain (7). These data suggest that while CorC and MgpA have roles in reducing excess-cation stress, these roles are distinct from that of PaeA, which is responsible for reducing cellular polyamine levels.

**Fig. 3.**
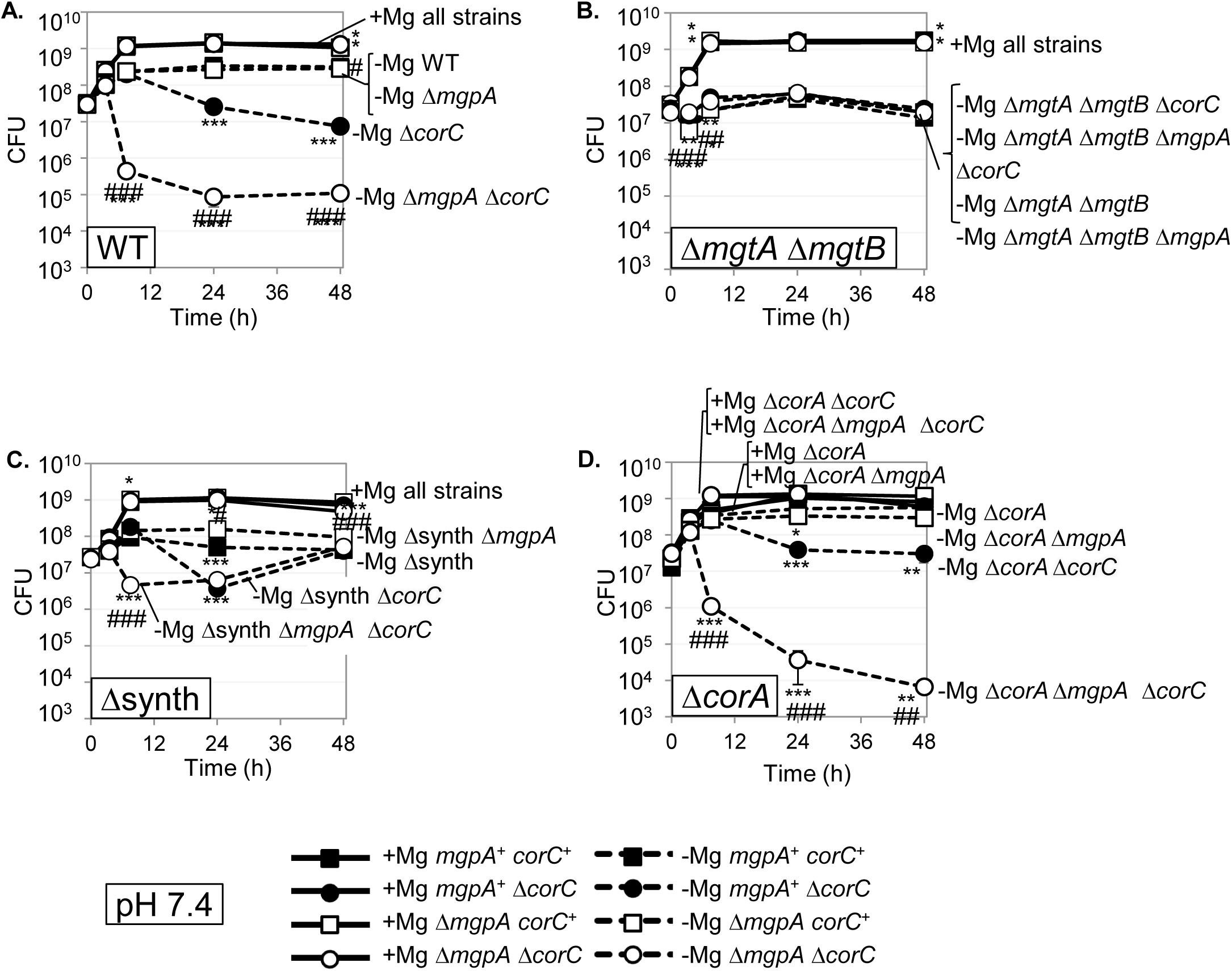
Loss of the high-affinity Mg²⁺ transporters MgtA and MgtB, but not CorA, suppresses the *corC* phenotype in stationary phase after Mg²⁺ starvation. The indicated strains were grown overnight in N-minimal medium pH 7.4 with 10 mM MgCl_2_, washed, diluted into N-minimal medium pH 7.4 with or without 10 mM MgCl_2_ (t=0 h), and incubated at 37℃. CFUs were determined at the indicated time points in WT (A), Δ*mgtA* Δ*mgtB* (B), Δsynth (C), and Δ*corA* (D) background. Values are mean ± SD, n = 6. Unpaired t test (p < 0.05*, 0.005**, 0.0005***) versus the *mgpA+ corC+* at the same time point and at the same Mg²⁺ concentration, and (p < 0.05^#^, 0.005^##^, 0.0005^###^) Δ*mgpA corC+* versus Δ*mgpA* Δ*corC* at the same time point and at the same Mg²⁺ concentration. Strains used: 14028, JS2692, JS2693, JS2694, JS2713, JS2714, JS2715, JS2716, JS2560, JS2717, JS2718, JS2717, JS2720, JS2721, JS2722, and JS2723.

### CorA is not required for CorC to function

The *corC* gene was first identified through a screen for mutations conferring resistance to cobalt (15). In addition to *corC*, this screen identified *corB* and *corA*, the latter encoding the primary Mg^2+^ transporter in Mg-replete conditions, capable of both Mg^2+^ import and efflux (24). Gibson et al. (15) showed that a *corBCD* triple mutant, similar to the *corA* mutant, lost all apparent Mg^2+^ efflux activity. To test if CorA is involved in the *corC* phenotype conferred in stationary phase after Mg^2+^ starvation, we examined the effect of deleting *corA* in the wild-type, Δ*corC*, Δ*mgpA*, and Δ*mgpA* Δ*corC* backgrounds. As shown in figure 3D, the loss of CorA had no significant effect on the Δ*corC* and Δ*mgpA* Δ*corC* phenotypes in stationary phase after Mg^2+^ starvation. Note that CorA does not function at low Mg^2+^ concentrations (25). Indeed, deletion of *corA*, *corB*, and *corD* had no effect on the *corC mgpA* phenotype (Fig. S4).

### Loss of CorC confers cobalt and manganese tolerance independent of MgpA

We also examined the role of CorC and MgpA in Co^2+^ and Mn^2+^ tolerance. As shown in figures 4A-C, deletion of *corC* increased tolerance to both Co^2+^ and Mn^2+^, similar to the phenotype conferred by deletion of *corA*, consistent with previous data (15). The double deletion of both *corC* and *corA*, which had not been examined in the previous study (15), exhibited a tendency to further increase tolerance to Co^2+^ or Mn^2+^ compared to the single deletion mutants. However, the degree of increase suggests that CorC and CorA act additively and independently. These results indicate that CorC does not require CorA to function. Interestingly, loss of MgpA had little effect on Co^2+^ or Mn^2+^ resistance in an otherwise wild type or *corC* background, but did enhance resistance in a *corA* background (Fig. 4A-C). Importantly, the Δ*paeA*, Δ*mgtA* Δ*mgtB*, Δsynth, and Δsynth Δ*mgtA* Δ*mgtB* mutants showed similar tolerance to Mn^2+^ as the wild type, further showing that the role(s) of CorC and MgpA is distinct from that of PaeA (Fig. 4D and E).

**Fig. 4.**
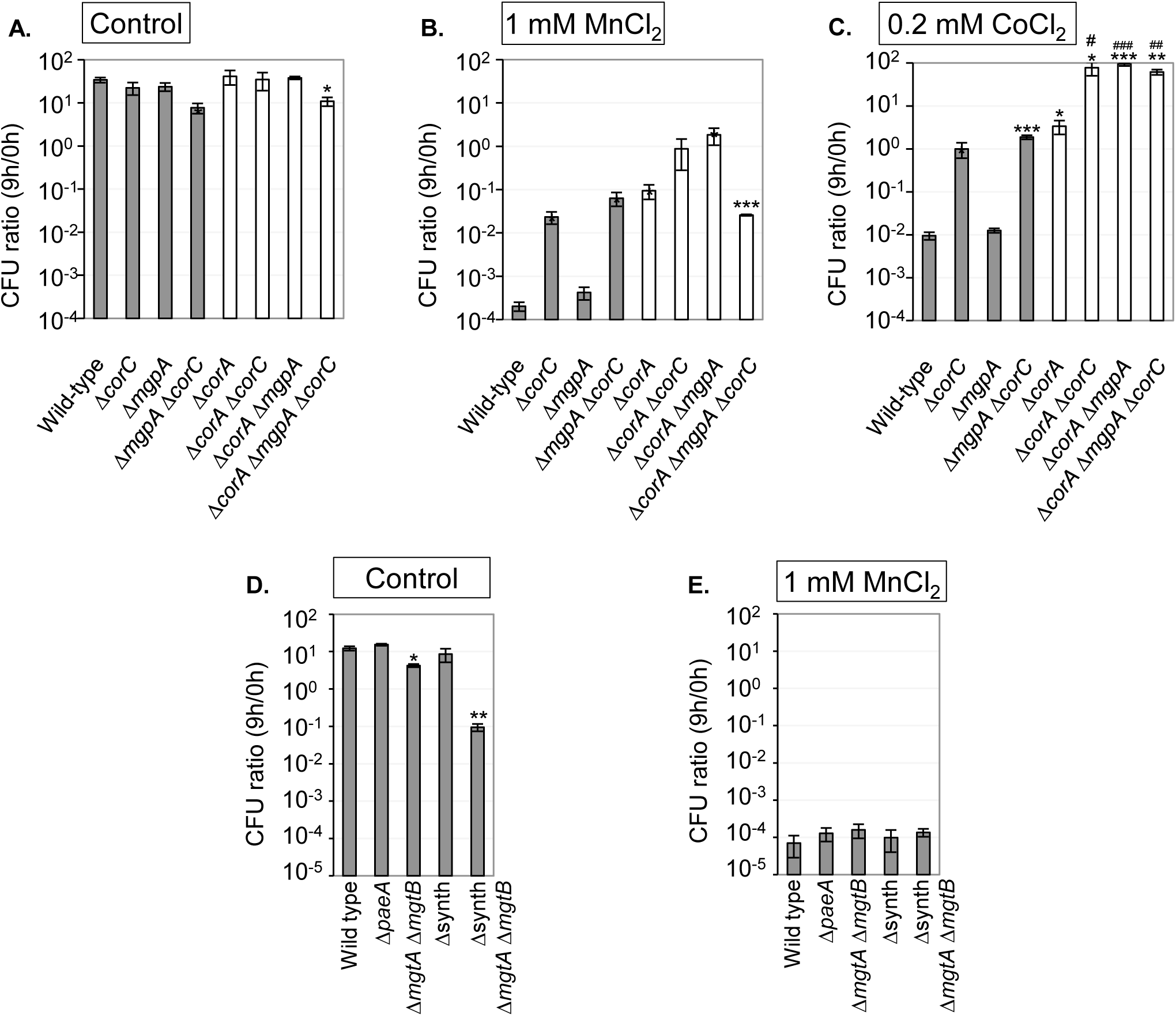
Loss of CorC, but not MgpA increases tolerance to Co^2+^ and Mn^2+^. The indicated strains were grown overnight in MOPS-minimal medium (pH 7.4) with 1.32 mM KH_2_PO_4_ and 10 mM MgCl_2_, washed, diluted into MOPS-minimal medium (pH 7.4) with or without the indicated amounts of MgCl_2_ or CoCl_2_ in the absence of KH_2_PO_4_ and MgCl_2_ (t=0h), and incubated at 37℃. CFUs were determined at 0h and 9h. CFU ratio (9h/0h) values are mean ± SD, n = 3. Unpaired t test (p < 0.05*, 0.005**, 0.0005***) versus WT and (p < 0.05^#^, 0.005^##^, 0.0005^###^) versus Δ*corA*. Strains used: 14028, JS2692, JS2693, JS2694, JS2720, JS2721, JS2722, JS2723, JS2430, JS2562, JS2560, and JS2563.

### ΔmgpA and ΔcorC mutants show higher tolerance to exogenous polyamines

The data above suggest that the CorC domain-containing proteins, CorC and MgpA, play a role in alleviating excess-cation stress after Mg^2+^ starvation, similar to the general role of PaeA. The specific role of PaeA is to efflux the divalent polyamines, cadaverine and putrescine, the synthesis of which is induced upon Mg^2+^ starvation (8, 14). Indeed, a *paeA* mutant is sensitive to exogenously added putrescine or cadaverine in stationary phase (14). To further examine the effect of the *corC* and *mgpA* genes on excess-cation stress, we measured tolerance to exogenous polyamines after growth in medium with high or low Mg^2+^. We tested their phenotype in a Δ*paeA* background because this strain is sensitive to polyamines and any additional effects should be easily detectable in this background. Strains were grown in medium with 50 μM MgCl_2_ (low Mg^2+^) or 10 mM MgCl_2_ (high Mg^2+^), washed, and incubated in a pH 8.5 buffer with the indicated polyamines. This level of low Mg^2+^ does not induce lethality in stationary phase. Note that high pH allows polyamines to be deprotonated and diffuse passively into the cell cytoplasm (14, 26).

As expected, the *paeA* mutant, grown to stationary phase in 10 mM Mg^2+^, is sensitive to either putrescine or cadaverine (Fig. 5). When cells were grown in low Mg^2+^, the sensitivity to polyamines was significantly reduced, as we have previously shown; it is the combined amount of Mg^2+^ and polyamine that is lethal (7). Deletion of both *corC* and *mgpA* did not further sensitize the *paeA* mutant to polyamines under the conditions we tested, but rather suppressed sensitivity. Deletion of either *corC* or *mgpA* alone had no effect when cells were grown in high Mg^2+^, but did suppress somewhat when cells were grown in low Mg^2+^ (Fig. 5). These results are consistent with the conclusion that, unlike PaeA, CorC and MgpA are not directly involved in reducing cytoplasmic polyamine levels.

**Fig. 5.**
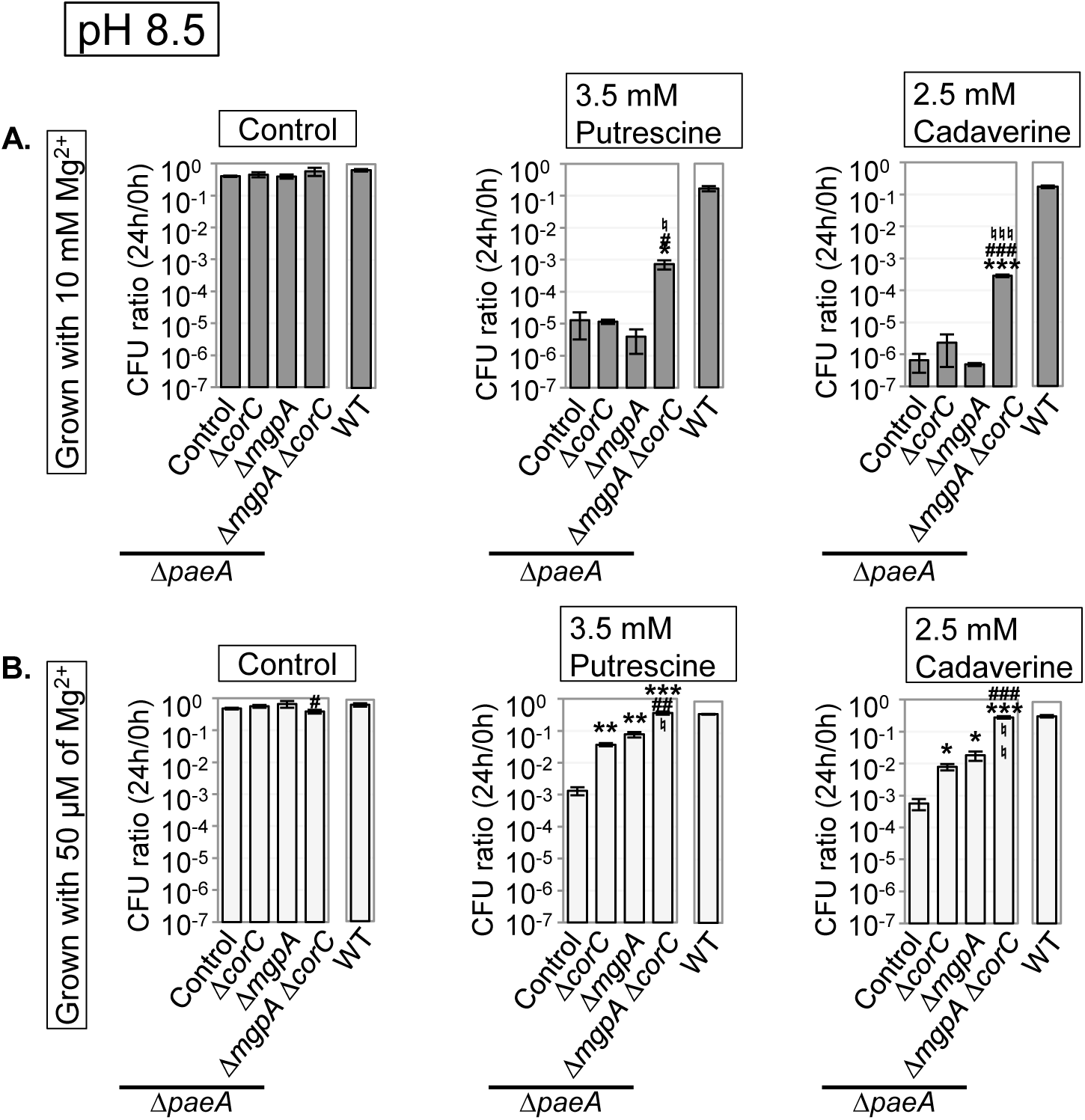
Loss of CorC and MgpA suppresses polyamine sensitivity in *paeA* background. The indicated strains were grown in N-minimal medium (pH 7.4) with (A) 10 mM or (B) 50 μM MgCl_2_ for 24 hours at 37℃, washed, diluted into pH 8.5-buffered saline containing indicated polyamines (t=0h), and incubated at 37℃. CFUs were determined at 0h and 24h. CFU ratio (24h/0h) values are mean ± SD, n = 3. Unpaired t test (p < 0.05*, 0.005**, 0.0005***) versus WT, (p < 0.05^#^, 0.005^##^, 0.0005^###^) Δ*mgpA corC+* versus Δ*mgpA* Δ*corC*, and (p < 0.05^♮^, 0.005 ^♮ ♮^, 0.0005 ^♮ ♮ ♮^) *mgpA+* Δ*corC* versus Δ*mgpA* Δ*corC*. Strains used: 14028, JS2695, JS2724, JS2725, and JS2726.

### MgpA and CorC are required for acute systemic infection and for egg white tolerance

We previously demonstrated that the polyamine efflux transporter PaeA is required for full virulence in mice (7). Interestingly, the virulence attenuation of *paeA* mutant varied depending on the mouse strain: C3H mice, with functional NRAMP1 (SLC11A1), versus BALB/c mice, which have a mutated NRAMP1 (27). NRAMP1 is a divalent cation transporter in the phagosomal membrane (28) that restricts *Salmonella* by reducing Mg²⁺ availability in phagosomes (29). The Δ*paeA* strain showed reduced virulence in both strains of mice, but the effect was stronger in the C3H background, supposedly due to lower Mg²⁺ levels in the phagosomes (7). To examine the effect of deleting the *mgpA* and *corC* genes in acute systemic infection *in vivo*, we performed competition assays after intraperitoneal infection (IP) in C3H mice or BALB/c mice. As shown in Table 1, the Δ*corC* Δ*mgpA* strain showed attenuated virulence only in BALB/c mice. In C3H mice, the Δ*mgpA* Δ*corC* strain was equally virulent to the wild type. Note that in the repeat of these experiments (second set of five mice each), the C3H and BALB/c mice were injected with the same inoculum. These results suggest that MgpA and CorC are important for virulence when the Mg²⁺ levels in the phagosome are only moderately limited.

**Table 1.** Competition assays with the Δ*mgpA* Δ*corC* mutant.

| Strain A<sup>a</sup> | Strain B<sup>a</sup> | Mouse Strain | No. of Mice | Organ | Median CI<sup>b</sup> | P value<sup>c</sup> | Fold change<sup>d</sup> |
| --- | --- | --- | --- | --- | --- | --- | --- |
| $\Delta mgpA$<br>$\Delta corC$ | WT | C3H | 10 | Spleen | 0.82 | NS | ~1 |
|  |  |  |  | Liver | 1.01 | 0.03 | ~1 |
|  |  | BALB/c | 10 | Spleen | 0.18 | <0.00005 | 5.6 |
|  |  |  |  | Liver | 0.10 | <0.00005 | 10.5 |

<sup>a</sup> Strains used were 14028 and JS2694.
<sup>b</sup> Bacteria were recovered from the spleen and liver after intraperitoneal (i.p.) competition assays. The competitive index (CI) was calculated as (percent strain A recovered/percent strain B recovered)/(percent strain A inoculated/percent strain B inoculated).
<sup>c</sup> Unpaired Student's t tests were used to compare the logarithmically transformed CI values to the inocula. NS, not significant.
<sup>d</sup> The fold attenuation is 1/CI, reflecting the level of attenuation of strain A relative to strain B.

To further investigate the role of CorC and MgpA in *Salmonella* pathogenesis, we next focused on egg white tolerance. It was reported that loss of MgpA (YoaE) confers sensitivity to egg white in *S*. Enteritidis, the major serovar linked to human salmonellosis from contaminated eggs (30). We tested egg white tolerance in wild type, Δ*mgpA*, Δ*corC*, and Δ*mgpA* Δ*corC* strains of *S*. Typhimurium. As shown in Figure 6A, deletion of both *mgpA* and *corC* conferred increased sensitivity to egg white, whereas the individual deletions had no effect. The introduction of plasmids encoding either CorC or MgpA suppressed this egg white sensitivity (Fig. 6B). These phenotypes differ from those seen in *S*. Enteritidis, in which the single deletion of the *mgpA* gene resulted in sensitivity to egg white (30). These results indicate that *Salmonella* Typhimurium requires either MgpA or CorC to protect from egg white.

**Fig. 6.**
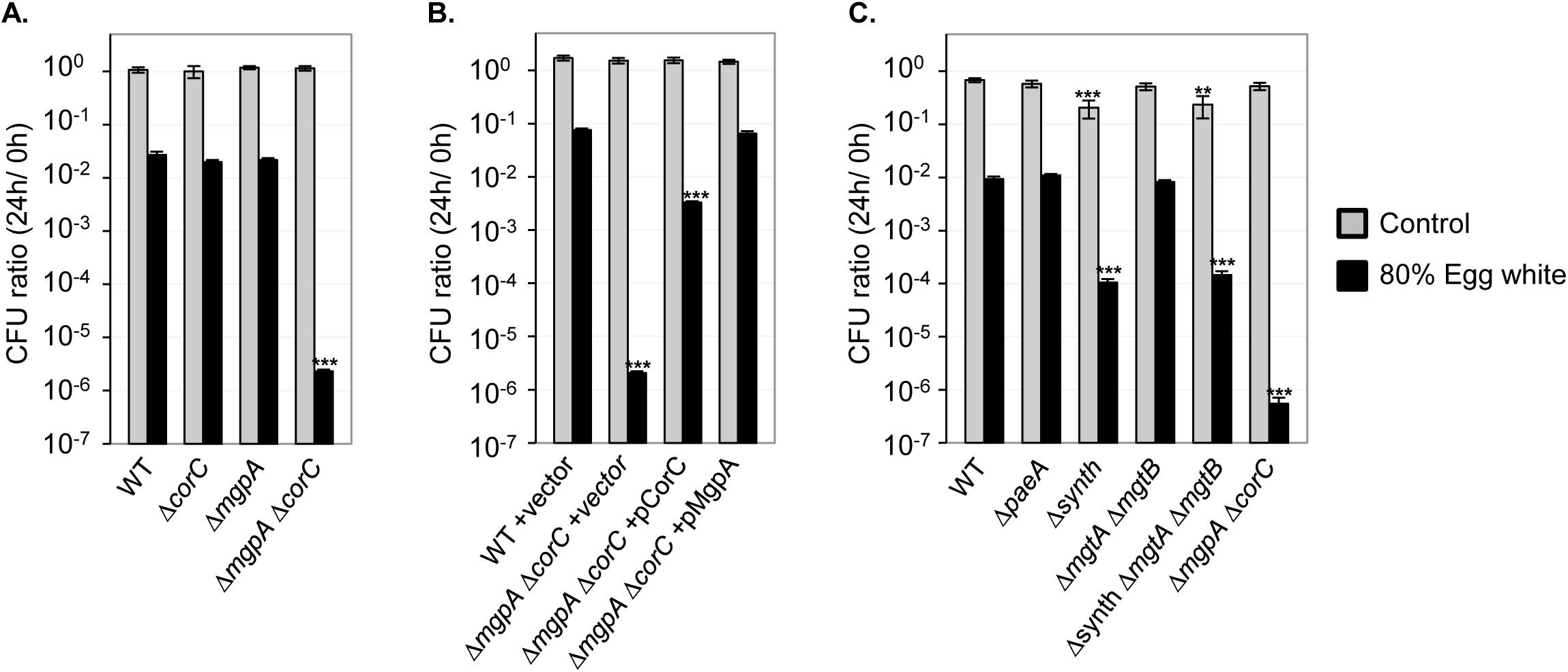
The Δ*mgpA* Δ*corC* strain is sensitive to egg white. The indicated strains were pre-grown to mid-exponential phase in N-minimal medium (pH 7.4) with 10 mM MgCl_2_ at 37℃, washed, diluted into 80% egg white solution or PBS (t=0 h), and incubated at 37℃. CFUs were determined at 0h and 24h. CFU ratio (24h/0h) values are mean ± SD, n = 6. Unpaired t test (*p* < 0.05*, 0.005**, 0.0005***) versus WT same treatment. Strains used: 14028, JS2728, JS2729, JS2730, JS2699, JS2731, JS2732, JS2733, JS2430, JS2560, JS2562, JS2563, and JS2694.

To understand the relationship between excess-cation stress and egg white stress, we examined whether other genes involved in reducing excess-cation stress are involved in egg white tolerance. As shown in Figure 6C, mutants lacking PaeA, responsible for polyamine efflux, or MgtA and MgtB, the inducible Mg^2+^ transporters, showed similar viability as the wild type upon egg white exposure, suggesting that these transporters are not involved in egg white tolerance. Interestingly, the Δsynth mutant, lacking genes for polyamine synthesis (7, 14), exhibited sensitivity to egg white, although the degree of viability loss in the Δsynth mutant was smaller than in the Δ*mgpA* Δ*corC* strain (Fig. 6C). These results suggest that polyamine synthesis is required for survival in egg white while polyamine efflux is not necessary. Altogether, we found that MgpA and CorC are required for egg white tolerance and for acute infection in the BALB/c mouse model. Conversely, PaeA is preferentially required for acute infection in the C3H mouse model, and has no apparent role during egg white tolerance. These findings show that the functions of MgpA and CorC are distinct from that of PaeA during *Salmonella* pathogenesis.

### MgpA and CorC phenotypes are affected by pH

Macrophages limit the growth of *Salmonella* not only by restricting Mg^2+^ availability (28, 29), but also by acidifying the phagocytic vacuole (31, 32). In contrast, egg whites are alkaline, with a pH that ranges from 7.6 when fresh to 9.2 as they age (33). The egg whites we used had a pH of about 8.6. This suggests that the phenotypes of the Δ*mgpA* Δ*corC* strains could be influenced by pH. We had tested the impact of deleting either or both the *mgpA* and *corC* genes on survival during stationary phase after Mg^2+^ starvation only at a pH of 7.4 (Fig. 2). To gain insight into the effect of pH on the phenotypes of the Δ*corC* and Δ*mgpA* Δ*corC* strains, we monitored the viability of wild-type, Δ*corC*, Δ*mgpA*, and Δ*mgpA* Δ*corC* strains after Mg^2+^ starvation when grown in acidic (pH 5.5) or alkaline (pH 8.5) N-medium. As shown in figure 7, when grown in acidic medium, the Δ*corC* strain survived after Mg^2+^ starvation for 48 hours at the same level as the wild-type. While the Δ*mgpA* Δ*corC* strain still exhibited a loss of viability in stationary phase when grown without Mg^2+^, the loss of viability was much reduced compared to that observed in neutral pH medium (Figs. 2C and 7A). In contrast, when grown in alkaline medium, both the Δ*corC* and Δ*mgpA* Δ*corC* strains exhibited a greater loss of viability in stationary phase after Mg^2+^ starvation compared to when grown in neutral pH medium (Figs. 2C and 7B). Altogether, the function of MgpA and CorC becomes more critical for survival in stationary phase after Mg^2+^ starvation as the pH rises.

**Fig. 7.**
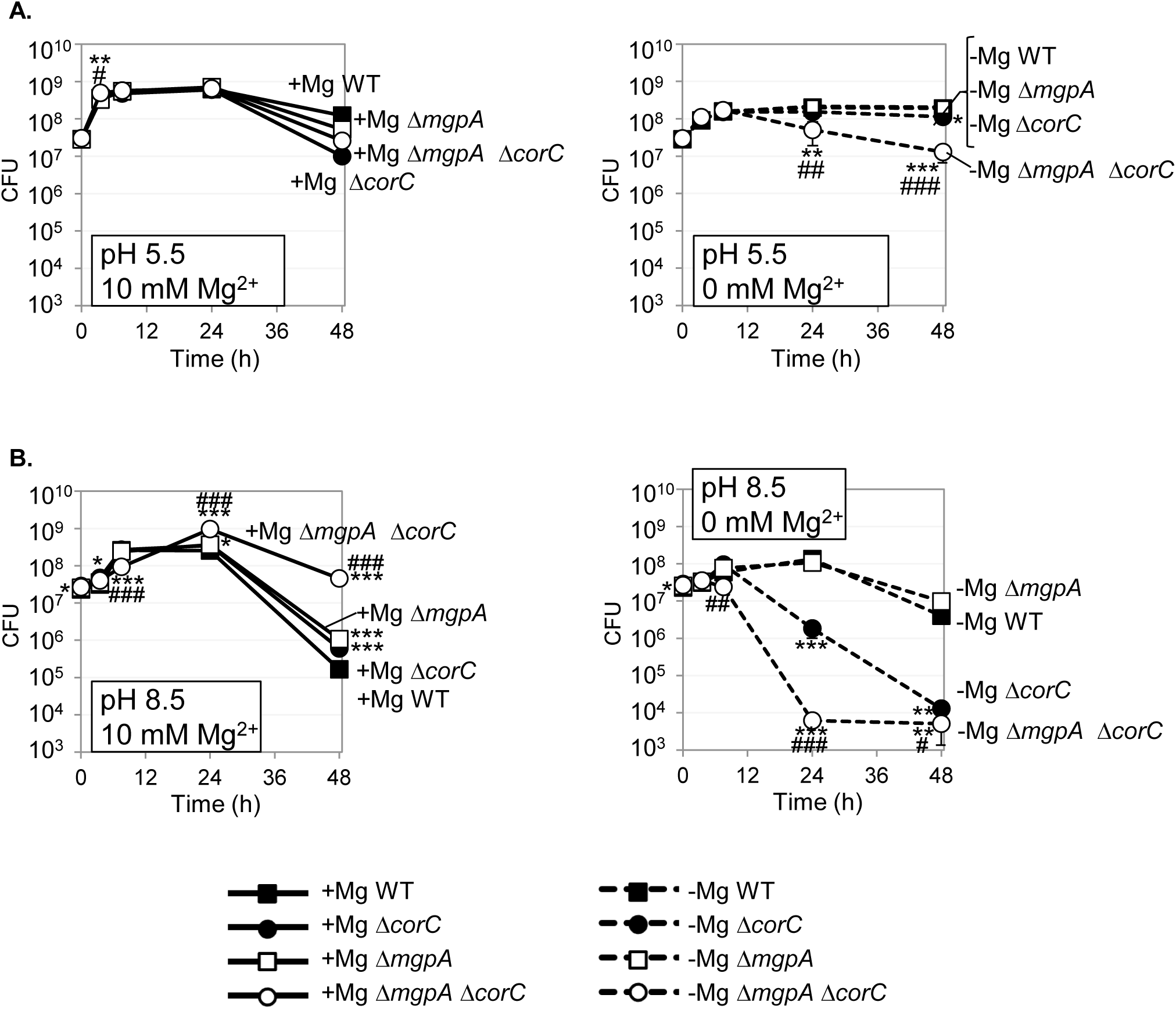
The loss of viability in Δ*corC* and Δ*mgpA* Δ*corC* strains during stationary phase after Mg²⁺ starvation is reduced in acidic medium and exacerbated in alkaline medium. The indicated strains were grown overnight in N-minimal medium pH 7.4 with 10 mM MgCl_2_, washed, diluted into N-minimal medium (A) pH 5.5 and (B) pH 8.5 with or without 10 mM MgCl_2_ (t=0 h), and incubated at 37℃. CFUs were determined at the indicated timepoints. Values are mean ± SD, n = 6. Unpaired t test (p < 0.05*, 0.005**, 0.0005***) versus corresponding WT and (p < 0.05^#^, 0.005^##^, 0.0005^###^) Δ*mgpA corC+* versus Δ*mgpA* Δ*corC* at the same time point and at the same Mg²⁺ concentration. Strains used: 14028, JS2692, JS2693, and JS2694.

### MgpA and CorC are required for growth and survival in high Mg^2+^ at alkaline pH

The data above show that MgpA and CorC are essential for survival during the stationary phase after Mg^2+^ starvation, particularly at neutral to alkaline pH. However, the Mg^2+^ concentration in egg white is reported to be 3.7-4.9 mM (34), which typically does not induce Mg^2+^ starvation in *Salmonella in vitro* (13). Therefore, we hypothesized that the Δ*mgpA* Δ*corC* strain is sensitive to Mg^2+^ in egg white at high pH. To test this hypothesis, the wild type, Δ*corC*, Δ*mgpA*, and Δ*mgpA* Δ*corC* strains were pregrown to the mid-exponential phase in pH 7.4 N-minimal medium with 1 mM MgCl_2_, then washed, diluted into pH 7.4 or pH 8.5 N-minimal medium with increasing amounts of MgCl_2_, and shaken vigorously at 37°C. OD_600_ and CFU were determined at the indicated time points.

When grown in pH 7.4 medium supplemented with 1, 10, or 100 mM MgCl_2_, all strains exhibited identical growth (Fig. 8A-C) and did not show any loss of viability (Fig. S5A-C). When grown in pH 8.5 medium supplemented with 1 mM MgCl₂, all strains exhibited identical growth (Fig. 8D) and did not lose viability (Fig. S5D). When grown in alkaline medium supplemented with 10 mM or 100 mM MgCl₂, the Δ*mgpA* Δ*corC* strain ceased growth after 3 hours of incubation (Fig. 8E and F) and, at 100 mM Mg^2+^, showed a drastic loss of viability after 12 hours of incubation (Fig. S5F). This phenotype is synthetic; the single deletion mutants showed only minor phenotypes. Collectively, these results show that the Δ*mgpA* Δ*corC* strain is sensitive to high Mg^2+^ in an alkaline medium.

**Fig. 8.**
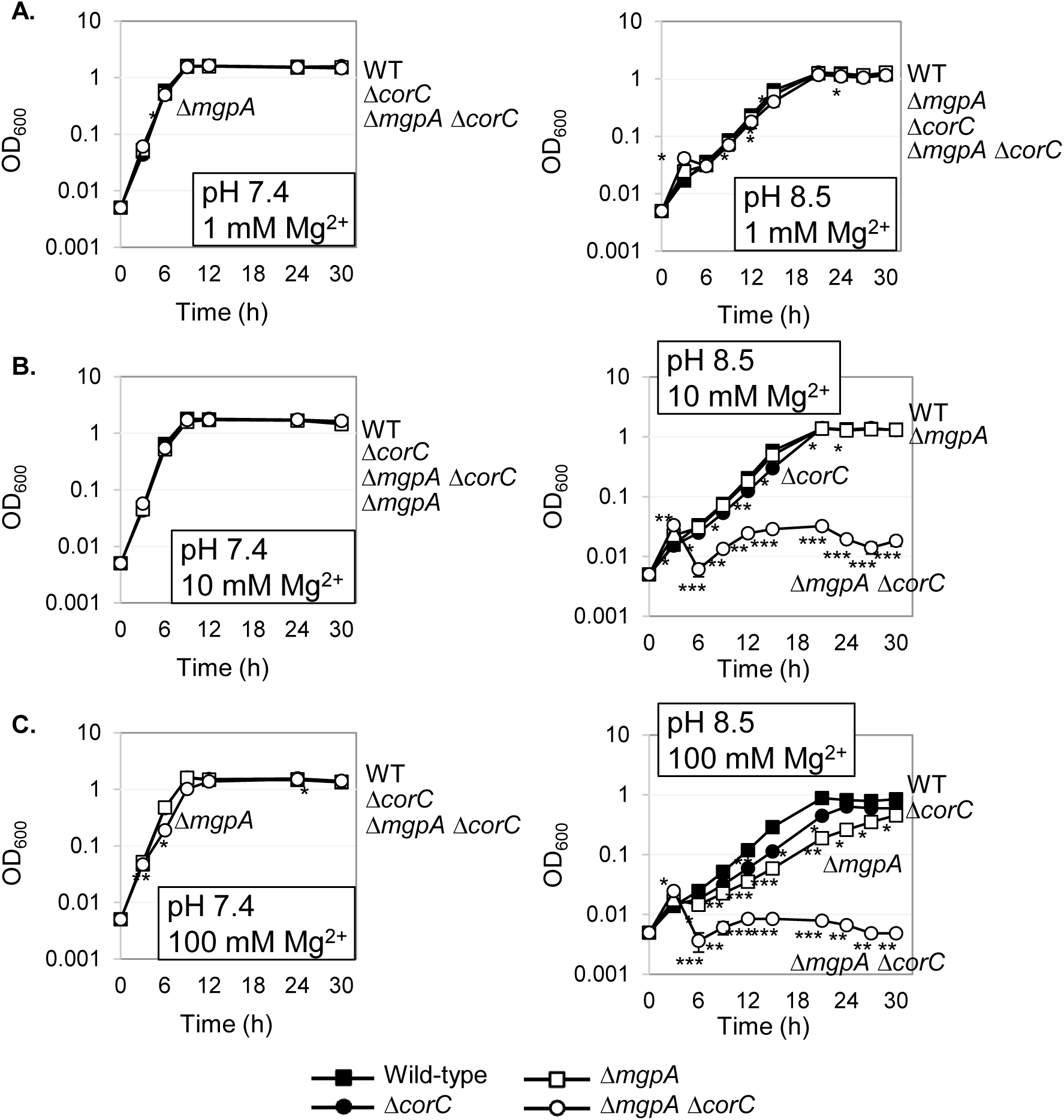
pH affects growth under high Mg2^+^ conditions. The indicated strains were pre-grown to mid-exponential phase in N-minimal medium pH 7.4 with 1 mM MgCl_2_, washed, and diluted into N-minimal medium pH 7.4 or pH 8.5 with (A) 1 mM, (B) 10 mM, and (C) 100 mM MgCl_2_ (t=0h), and incubated at 37℃. OD_600_ was determined at the indicated timepoints. Values are mean ± SD, n = 3. Unpaired t test (p < 0.05*, 0.005**, 0.0005***) versus corresponding WT at the same timepoint. Strains used: 14028, JS2692, JS2693, and JS2694.

We also tested the roles of polyamines and PaeA under these conditions. Importantly, the Δ*paeA* strain did not exhibit any growth defects or loss of viability under high Mg^2+^ stress at pH 7.4 or 8.5 (Fig. S6), consistent with CorC and MgpA having functions distinct from that of PaeA. Interestingly, the strain incapable of synthesizing polyamines (Δsynth) showed a severe growth defect at pH 8.5 compared to pH7.4 (Fig S6), but no loss of viability (Fig. S7). This defect was independent of Mg^2+^ concentration.

### The *corC* and *mgpA* genes are regulated in response to high Mg^2+^ and pH

We examined the expression of *corC* and *mgpA* in *Salmonella* using *lacZ* translational fusions. These constructs (Fig. 9A), containing the regions upstream of *corC* or *mgpA* and part of the open reading frames fused in frame to *lacZ*, were inserted at the λ *att* site in the chromosome. Thus, the chromosomal *corC* and *mgpA* loci were not disrupted. We measured expression after growth in different pH conditions (acidic [pH 5.5], neutral [pH 7.4], or alkaline [pH 8.5]) and in varying Mg^2+^ concentrations (0, 0.05, or 10 mM MgCl_2_). Figure 9B shows that *corC* expression increased under high Mg^2+^ (10 mM) conditions, regardless of the pH of the medium. Expression of *mgpA* was elevated in high Mg^2+^ (10 mM) when the pH was neutral or alkaline (Fig. 9C). Thus, expression of both CorC and MgpA are induced by high Mg^2+^ concentrations, consistent with their proposed role in combating excess-cation stress, with phenotypes exacerbated at high pH.

**Fig 9.**
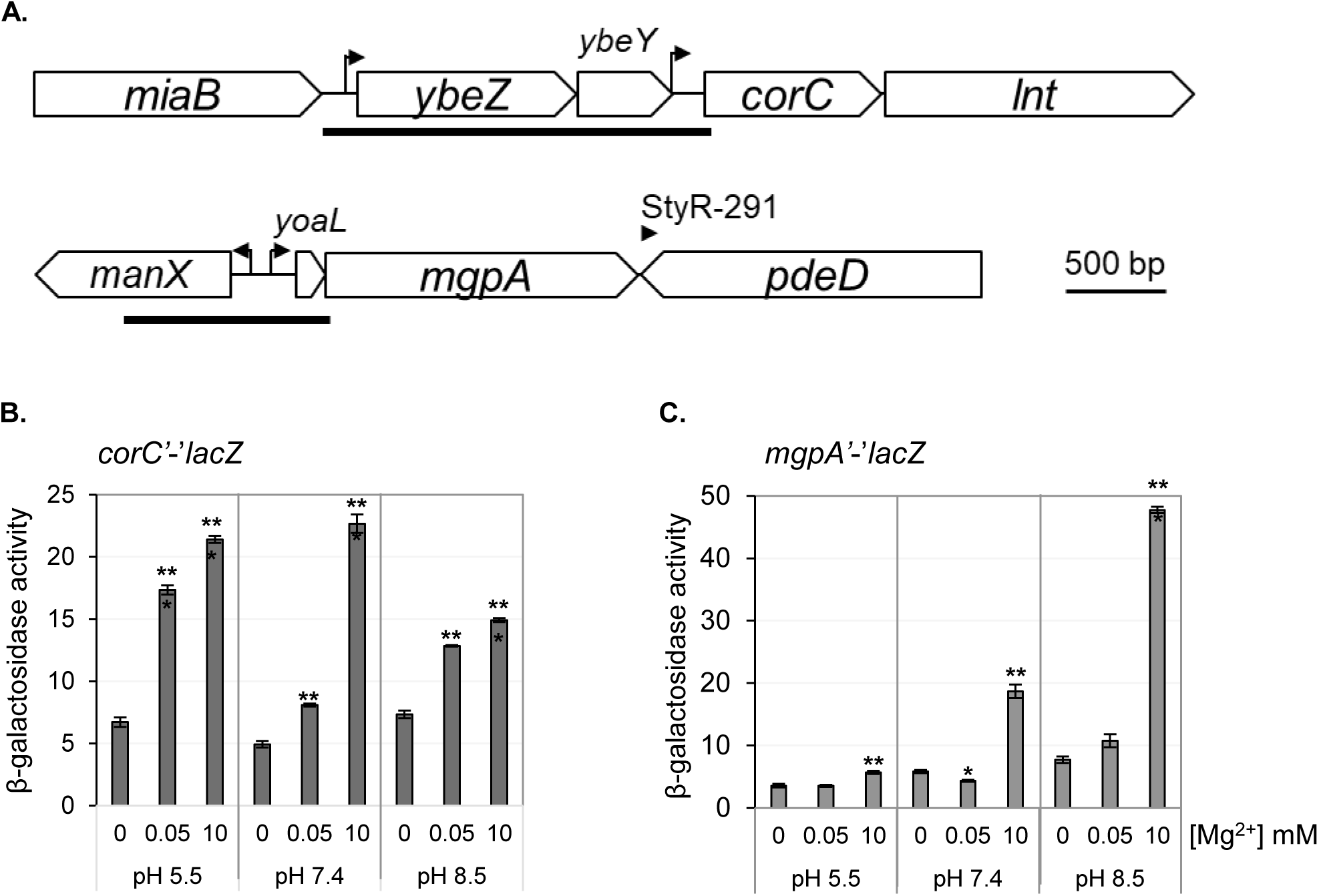
Expression of *mgpA* and *corC* are upregulated under high Mg^2+^ conditions. (A) The *corC* and *mgpA* loci. Dark bars indicate the sequences cloned upstream and in-frame with *lacZ*. The (B) *corC’-’lacZ* and (C) *mgpA’-’lacZ* fusion strains were pre-grown to mid-exponential phase in N-minimal medium pH 7.4 with 1 mM MgCl_2_ at 37℃, washed, diluted in N-minimal medium pH 5.5, pH 7.4, and pH 8.5 with or without the indicated amount of MgCl_2_, and incubated at 37℃ for 24h. β-galactosidase activity was measured as described in Materials and Methods section. Values are mean ± SD, n = 6. Paired t test (*p* < 0.05*, 0.005**, 0.0005***) versus 0 mM Mg^2+^ at the same pH. Strains used: JS2734 and JS2735.

### Absolute Mg levels are not significantly altered in the *corC* and *mgpA* mutants

The phenotypes conferred by loss of MgpA and CorC are clearly affected by Mg^2+^ levels in the media. A simple model would suggest that MgpA and CorC either directly or indirectly affect intracellular Mg^2+^ levels. We measured intracellular Mg^2+^ levels using two different methods. Total cellular Mg^2+^ was assayed using inductively coupled plasma mass spectrometry (ICP-MS). Cells were grown to early stationary phase (7.5 hr) as in figure 2. The results indicated that total intracellular Mg^2+^ concentrations were approximately equal in all the various mutants, despite the fact that the *corC mgpA* double mutant had started to lose viability at this time point (Fig. S8).

To measure “free” Mg^2+^, we used a fluorescent Mg^2+^ sensor, MagFET (35). We first confirmed that fluorescence in the MagFRET system was directly proportional to Mg^2+^ concentrations in permeabilized cells (Fig. S9). We are, however, reluctant to draw any conclusions regarding absolute levels of available Mg^2+^ using this system. The wild type, Δ*corC*, Δ*mgpA*, and Δ*mgpA* Δ*corC* strains were grown in N-minimal medium at pH 7.4 with various concentrations of Mg^2+^ and samples were taken at 3 hr, 7.5 hr, and 24 hr. The OD_600_ of each sample was determined (Fig. S10) and then adjusted so that approximately equal numbers of cells were used to measure fluorescence. The CFUs present in each adjusted sample were also determined to measure viability of cells (Fig. S10). Interestingly, in growing cells, free Mg^2+^ concentrations apparently decreased as the Mg^2+^ in the medium increased, suggesting clear regulation of cations in response to the environment (Fig. 10). While total Mg^2+^ concentrations were unchanged in the Δ*mgpA* Δ*corC* double mutant, free Mg^2+^ concentrations were significantly lower at higher external concentrations at both 7.5 hr and 24 hr. Although counterintuitive, loss of CorC and MgpA are clearly affecting cation concentrations or cation distribution in the cell in response to external Mg^2+^ concentrations.

**Fig 10.**
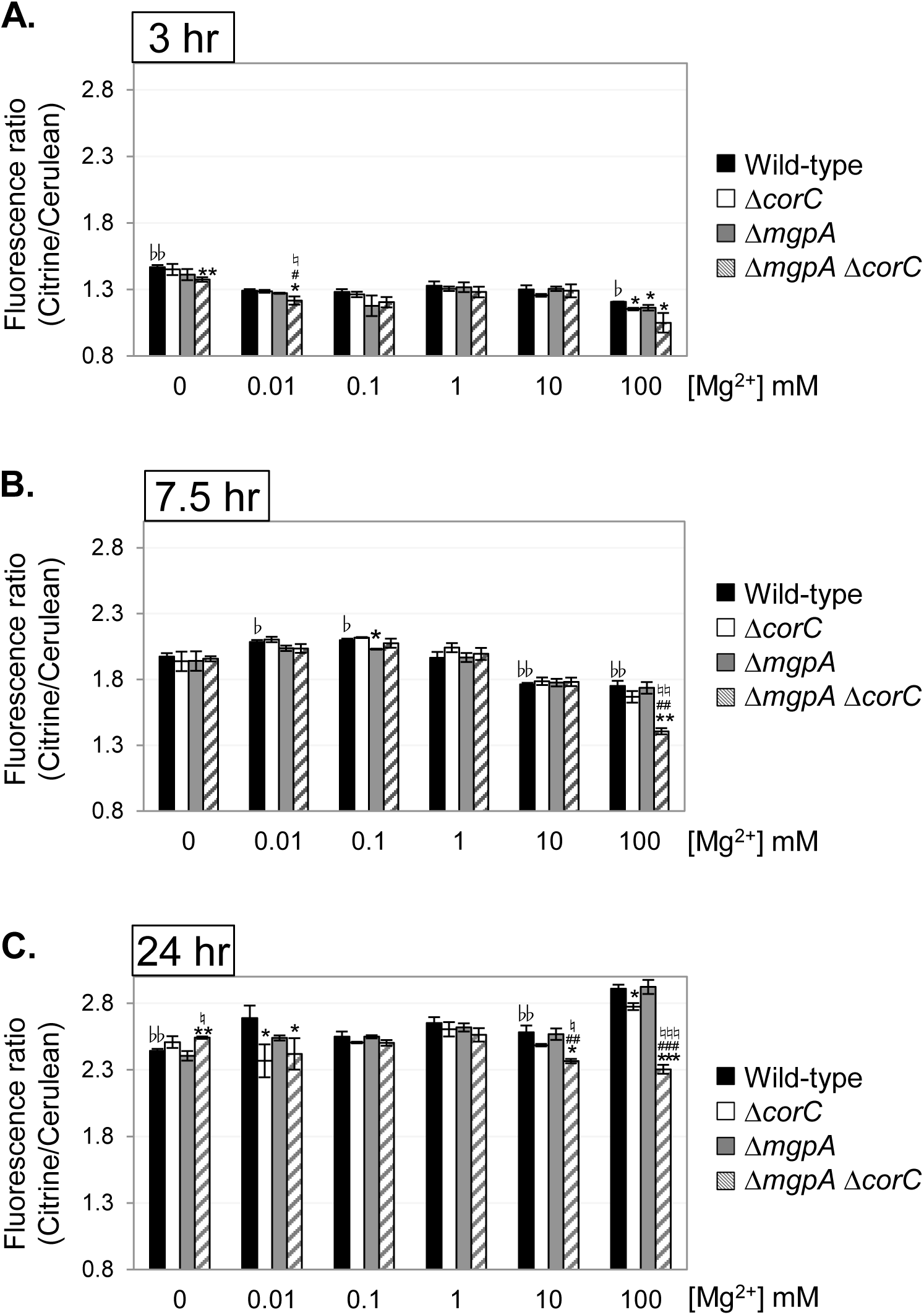
The Δ*corC* strain shows a lower free-Mg level at stationary phase when grown with 100 mM MgCl_2_, and the deletion of *mgpA* exacerbates the effect. The indicated strains harboring pS2513-MagFRET, pre-grown to mid-exponential phase in N-minimal medium (pH 7.4) with 1 mM MgCl_2_, were washed and diluted into N-minimal medium pH 7.4 with the indicated amounts of MgCl_2_, and incubated at 37 ℃. Aliquots (1.5 mL) were collected after (A) 3, (B) 7.5, and (C) 24 hours, and the fluorescence ratio (Citrine/Cerulean) was determined at OD₆₀₀ = 0.1. A higher ratio indicates a higher free-Mg level. Values are mean ± SD, n = 3. Unpaired t test (P < 0.05*; P < 0.005**; P < 0.0005***) versus the wild-type, (P < 0.05#; P < 0.005##; P < 0.0005###) of *mgpA* ^+^ Δ*corC* versus Δ*mgpA* Δ*corC*, or (P < 0.05♮; P < 0.005 ♮ ♮; P < 0.0005 ♮ ♮ ♮) of Δ*mgpA corC ^+^* versus Δ*mgpA* Δ*corC* at the same condition. Paired t test (P < 0.05♭; P < 0.005♭♭; P < 0.0005♭♭♭) versus 1 mM Mg^2+^ in the wild-type. Strains used: JS2736, JS2737, JS2738, and JS2739.

## DISCUSSION

Maintaining proper cation homeostasis in response to changes in the environment is essential for all living organisms. Previous studies reveal that, upon Mg^2+^ starvation, *Salmonella* induces the Mg^2+^ transporters MgtA and MgtB (9–13) and stimulates the synthesis of polyamines: cadaverine, putrescine, and spermidine (7, 8). Although the specific roles of polyamines are not fully understood, we propose that they act as simple cations that can substitute for Mg^2+^ to neutralize negative charges within cells. Once the Mg^2+^ concentration is re-established, the inner membrane transporter PaeA is required to efflux putrescine and cadaverine from the cell. Otherwise, when *Salmonella* enters stationary phase, the cell experiences what we term “excess-cation stress” (7). In the Δ*paeA* strain, polyamine accumulation occurs, leading to a loss of viability after Mg^2+^ starvation in stationary phase. This loss of viability can be suppressed by deleting the polyamine synthesis genes or the inducible Mg^2+^ transporter genes, showing that it is the combined concentration of polyamines and Mg that is lethal in stationary phase (7).

PaeA contains a CorC domain at its C-terminus. Although some CorC domain-containing proteins have been implicated in metal ion homeostasis, the exact function of this domain is not understood. Here we have shown that two additional CorC domain-containing proteins in *Salmonella*, CorC and MgpA (YoaE), play a role in protecting against excess-cation stress in stationary phase. After growth in Mg^2+^-limiting medium, a *corC* deletion mutant loses viability in stationary phase. Deletion of *mgpA* alone has no effect but confers a synthetic phenotype in the *corC* background; the double mutant suffers a precipitous loss of viability in stationary phase after Mg^2+^ starvation. These proteins act independently of PaeA. Deletion of *paeA* has an apparent additive effect in the *corC* and *corC mgpA* backgrounds, and deletion of *mgpA* confers no additional phenotype in the *paeA* mutant. Loss of viability in the *corC mgpA* double mutant is completely suppressed by loss of the inducible Mg^2+^ transporters, MgtA and MgtB, and partially suppressed by deletion of the polyamine synthesis genes. Thus, CorC and MgpA carry out some seemingly redundant function that is distinct from PaeA, yet all act to prevent excess-cation stress in stationary phase.

The concentrations of both Mg^2+^ and polyamines in *Salmonella* range from 5 - 40 mM and are inversely and coordinately regulated by unknown mechanisms (7). Both Mg^2+^ and polyamines have high affinities for phosphate-containing compounds and should compete for binding. For example, putrescine has an ∼10-fold higher affinity for ATP than does Mg^2+^ (36). It is also known that putrescine can functionally replace ∼80% of the Mg^2+^ in the ribosome (37). The cause of lethality under our conditions is not clear. Simplistically, lethality occurs when the combined concentrations of Mg^2+^ and polyamines are too high. However, it seems more likely that it is competition for binding to some specific compound in the cell that is interfering with an essential function(s). Thus, it is the balance of cations and binding sites that is critical, rather than absolute levels. Our data are all consistent with PaeA exporting putrescine and cadaverine (7, 14). Previous data implicate CorC in Mg^2+^ efflux (15). Given the apparent redundancy between MgpA and CorC, as evidenced by the synthetic phenotypes, we presume that MgpA is also an ion transporter. However, given our overall results, we cannot conclude that CorC or MgpA are directly involved in Mg^2+^ transport. Rather, they could control some other ion that is competing with, or for, Mg^2+^ and polyamines.

Further evidence shows that CorC/MgpA are functionally distinct from PaeA. The Δ*paeA* mutant, grown to stationary phase and suspended in high pH buffer, is sensitive to added cadaverine or putrescine (14). The polyamines are deprotonated at high pH, passively enter the cells, and, in the absence of PaeA, build to lethal concentrations. This lethality can be moderated by pre-growing the cells in low Mg^2+^ (7). Loss of CorC and MgpA partially suppresses this sensitivity to polyamines. In this situation, where polyamines continue to enter the cytoplasm, the functions of CorC and MgpA negatively affect survival. One possibility is that the increased levels of polyamines compete with Mg^2+^, and this liberated Mg^2+^ results in a shift in ion balance and loss of viability. In trying to compensate for this stress, CorC and MgpA are contributing to cation imbalance in this unusual situation.

Loss of CorC confers resistance to a high concentration of Mn^2+^ or Co^2+^ in the medium. MgpA has a more minor role in this phenotype. Loss of CorA also confers resistance to Mn^2+^ or Co^2+^, as previously shown (15). Deletion of *corC* in the *corA* background confers additional protection. CorA also did not affect lethality in stationary phase after Mg^2+^ starvation. Thus, CorC acts independently of CorA. Indeed, none of the previous cobalt-resistance loci, *corA*, *corB*, or *corD* (15), affect lethality in stationary phase after Mg^2+^ starvation. When extracellular Mn^2+^ or Co^2+^ enter the cytoplasm, the functions of CorC and MgpA negatively impact cell survival. These results suggest that, analogous to sensitivity to external polyamines, when Mn^2+^ or Co^2+^ enter the cytoplasm, primarily through CorA-dependent mechanisms (38, 39), the increased levels of Mn^2+^ or Co^2+^ compete with and liberate Mg^2+^. CorC and MgpA are somehow adjusting ion concentrations to compensate, but this results in increased toxicity and a loss of viability (Fig. 4A-C).

The Δ*corC* Δ*mgpA* mutant also behaves very differently than the Δ*paeA* mutant in two virulence models. The Δ*corC* Δ*mgpA* double mutant is sensitive to egg white, whereas loss of PaeA has no apparent effect. Huang et al. (30) demonstrated that the combination of high pH and a small peptide (smaller than 3 kDa) that is sensitive to proteinase K is necessary for egg white to kill the *mgpA* (*yoaE*) strain in *S*. Enteritidis. Further investigation into the molecular mechanisms of MgpA and CorC will deepen our understanding of how egg white kills *Salmonella*.

The Δ*corC* Δ*mgpA* mutant is also attenuated in competition assays in BALB/c mice, but, strikingly, there is no phenotype in C3H mice (Table 1). In contrast, the Δ*paeA* mutant shows a stronger attenuation in C3H mice, although there is still some phenotype in the BALB/c background (7). NRAMP1 is a divalent cation transporter in the phagosomal membrane that restricts *Salmonella* by reducing Mg²⁺ availability in phagosomes (28, 29). C3H mice have a functional NRAMP1 (SLC11A1), while BALB/c mice have a mutated NRAMP1 (27). It is clear that the macrophage phagosome in BALB/c is low in Mg²⁺ because the PhoPQ operon is strongly induced (7, 40). Presumably, the Mg²⁺ concentration is higher in the BALB/c phagosome than in the C3H phagosome, which could explain the increased sensitivity of the Δ*corC* Δ*mgpA* mutant. However, we cannot rule out other parameters in the phagosome that could differ between the mouse strains, affecting the overall environment and/or *Salmonella*’s regulatory response.

The pH of the media affects the phenotypes. The *corC mgpA* mutant loses viability after growth in Mg^2+^ starvation conditions in pH 7.4 or 8.5 medium. However, this phenotype is dramatically reduced when the pH of the medium is 5.5. High pH also enhances sensitivity to high Mg²⁺ in the medium; the *corC mgpA* mutant cannot grow in pH 8.5 medium when the Mg^2+^ concentration is ≥ 10 mM, whereas the mutant grows fine in pH 7.5 medium, even at 100 mM Mg^2+^. There are several possibilities for these phenotypic differences at acidic, neutral, and alkaline pH. One possibility is that there are one or more redundant systems that reduce excess-cation stress in acidic environments. For example, PaeA and the cadaverine exporter CadB are redundant in acidic conditions (7). Another possibility is that the stress itself is reduced or alleviated in acidic environments, such that CorC and MgpA are no longer required. High pH in the medium can also significantly affect the activity of bacterial transporters, particularly those that rely on proton-motive force (41). Thus, under high pH conditions, bacteria might be particularly sensitive to high or low cation concentrations in the medium. However, these phenotypes are seemingly contradictory to the mouse phenotypes, given that the macrophage phagosome is low pH. Further investigation is necessary to address these questions.

Consistent with their roles in protecting against excess-cation stress, we demonstrated that *mgpA* expression is elevated in high Mg^2+^ (10 mM) when the pH is neutral or alkaline (Fig. 9C), whereas *corC* expression increases under high Mg^2+^ (10 mM) conditions, regardless of the pH of medium (Fig. 9B). These results support the model that CorC and MgpA are involved in mitigating excess-cation stress, specifically high Mg^2+^ stress. The regulatory mechanisms governing *mgpA* and *corC* expression remain unclear. Chadani et al. showed that the translation of the N-terminal region of CorC (YbeX) contains an intrinsic ribosome destabilization (IRD) sequence (42). This IRD sequence destabilizes the translating ribosomal complex, leading to premature termination of translation under conditions in which ribosome function is reduced. Low Mg^2+^ stress is a well-known example of ribosome-destabilizing conditions. For example, under such stress, the translation of MgtL, a leader peptide for the inducible magnesium transporter MgtA, stalls due to an IRD mechanism, resulting in the upregulation of *mgtA* expression (43, 44). Consequently, CorC expression is likely upregulated when cytoplasmic Mg^2+^ levels exceed a certain threshold. As for *mgpA* regulation, a leader peptide, YoaL, overlaps with the upstream region of the *mgpA* gene (Fig. 9A; (45). Replacing the start codon of YoaL with a stop codon leads to decreased MgpA translation (46). This suggests that YoaL might play a role in upregulating *mgpA* expression in response to high Mg^2+^ levels. Further research is required to test this hypothesis and to identify the signals that influence this leader peptide-mediated regulation of *mgpA* expression.

Based on the presence of a transmembrane TerC-domain in the N-terminus of MgpA (**Fig. 1**), which is similar to a protein in *Bacillus subtilis* known to be involved in Mn transport (47), MgpA appears to function directly as a cation transporter. In contrast, CorC, unlike other CorC domain-containing proteins, is cytoplasmic and contains only a CBS-pair domain and the CorC domain (**Fig. 1**). Studies have shown that inactivation of the *corC* gene results in decreased Mg^2+^ efflux activity under high Mg^2+^ conditions. When *corC* is inactivated along with *corB* and *corD* (*apaG*) genes, Mg^2+^ efflux activity is completely lost (15). These data would suggest that CorC regulates a protein responsible for Mg^2+^ efflux activity, although we cannot rule out regulation of some other ion that is competing for Mg^2+^. However, the protein that CorC regulates does not appear to be CorA, CorB, CorD, or any other CorC-domain-containing proteins, including MgpA, because deletion of the *corC* gene in strains lacking these proteins still confers phenotypes. Therefore, we hypothesize that CorC mitigates excess-cation stress, in response to Mg^2+^ levels, by regulating an uncharacterized transporter. This also leads to the general conclusion that the CorC domain, likely in conjunction with the CBS domain, is regulatory, perhaps sensing levels of cytoplasmic cations. Indeed, a crystal structure of the CorB CBS-pair from the thermophilic archaeon *Methanoculleus thermophilus* shows that dimerization is dependent on binding of ATP-Mg^2+^, with the Mg^2+^ ion coordinated between the dimers (16), potentially providing a mechanism for sensing relative Mg^2+^ levels.

Mg^2+^ is critical for ribosome assembly, function, and stability (4, 6). It is also required for stability of the outer membrane, neutralizing the negative charges in outer membrane lipopolysaccharide (LPS) (5, 48). Indeed, low Mg^2+^ induces stress on outer membrane homeostasis and biogenesis (49). Interestingly, the *corC* gene is in an operon downstream of *ybeZ* and *ybeY*, and upstream of *lnt*. In addition to the promoter upstream of *ybeZ*, there is a start site of transcription between *ybeY* and *corC* (Fig 9; (21, 50)). YbeY is an endoribonuclease with pleiotropic effects on ribosome maturation and function (51). The upstream YbeZ is a putative RNA helicase that is also proposed to participate in ribosome maturation (52). Lnt (CutE), encoded downstream of CorC, is an essential protein required for the final acylation of the N-terminal Cys residue of processed outer membrane lipoproteins (53), including the major lipoprotein Lpp, which tethers the outer membrane to the peptidoglycan (54).

Sarigul et al. (22) explored the role of CorC in ribosome metabolism in *E. coli*. They demonstrated that a *corC* deletion mutant exhibited an extended lag phase coming out of stationary phase, particularly when cells were grown at elevated temperatures and under low Mg^2+^ conditions. The *corC* mutant also displayed increased sensitivity to several ribosome-targeting antibiotics. The accumulation of 17S pre-rRNA, a precursor of 16S rRNA, and partial degradation intermediates of 16S rRNA were noted in the *corC* deletion strain upon entering stationary phase. Importantly, the growth phenotypes were suppressed by adding additional Mg^2+^ to the growth medium. Orelle et al. (55) constructed an *E. coli* “Ribo-T” strain that expresses an engineered hybrid rRNA composed of both 16S and 23S rRNA such that the ribosome subunits are tethered and inseparable. A strain making only these engineered ribosomes grew slowly. Selecting for a faster growing mutant yielded a nonsense mutation in *corC,* together with a missense mutation in *rpsA*. Thus, loss of CorC affects ribosome stability and/or function in a Mg^2+^ dependent manner. Overall, the coordinate regulation of these various genes suggests that CorC has an important role in both Mg^2+^-dependent ribosome function and outer membrane stability. Further investigation into the biochemical function of CorC and MgpA is required to understand how these proteins reduce excess-cation stress after Mg^2+^ starvation and how regulation of cation levels affects overall cell physiology and survival in host tissues.

## EXPERIMENTAL PROCEDURES

### Media

During construction of strains, bacteria were routinely grown at 37°C in Luria-Bertani (LB) medium containing 1% NaCl. Strains containing temperature-sensitive plasmids pCP20 and pKD46 were grown at 30°C. When necessary, antibiotics were used at the following concentrations: ampicillin, 50 μg/ml; chloramphenicol, 10 μg/ml; kanamycin, 50 μg/ml; apramycin, 50 μg/ml. N-minimal medium (9, 56) contains 5 mM KCl, 7.5 mM (NH_4_)_2_SO_4_, 0.5 mM K_2_SO_4_, 1 mM KH_2_PO_4_, 0.1 M Tris, 15 mM glycerol, and 0.1% casamino acid. The pH was adjusted as indicated. MOPS medium (57), adjusted to pH 7.4 with KOH, contains 40 mM MOPS, 4 mM Tricine, 9.5 mM NH₄Cl, 0.276 mM K₂SO₄, 50 mM NaCl, and 0.1 % casamino acids. When indicated, 1.32 mM K₂HPO₄ and 10 mM MgCl₂ were added. We did not add the trace metals listed in the original recipe (57) to minimize the effects of metal addition.

### Bacterial strains and plasmids

Strains are described in Table S1, while plasmids are listed in Table S2. All *Salmonella* strains used are derivatives of *Salmonella enterica* serovar Typhimurium strain 14028. Deletions with concomitant insertion of antibiotic resistance cassettes were constructed using λ Red-mediated recombination as previously described (58, 59), with the indicated endpoints (Table S1). Deletions were verified by PCR analysis, and then transduced into the approrpiate backgrounds using phage P22 HT105/1 int-201 (60). All plasmids were passaged through a restriction-minus modification-plus Pi^+^ *Salmonella* strain (JS198) (59) prior to transformation into *Salmonella* strains. Antibiotic resistance cassettes were removed using the pCP20 plasmid. The resulting deletions were confirmed by PCR.

The translational out of locus *corC’-‘lacZ* and *mgpA’-‘lacZ* translational fusions were constructed by introducing the regulatory regions of *corC* and *mgpA*, along with the first or twentieth amino acids of the ORFs, respectively, amplified by PCR using primers listed in Table S3, upstream and in-frame with the promotorless *lacZ* in the pDX1 vector via Gibson assembly (NEBuilder® HiFi DNA Assembly)(Fig 9). The resulting product was transformed into DH5α λ*pir*, and the plasmid was confirmed by DNA sequencing. The plasmids, pDX1-*corC-lacZ* and pDX1-*mgpA-lacZ*, were integrated into the λ-attachment site in the *Salmonella* chromosome, as previously described (61). The primers used and the endpoints of the cloned fragments are indicated in Tables S3 and S2, respectively.

To construct the pS2513-MagFRET plasmid, the ratiometric fluorescent indicator protein PHP in the pS2513-PHP plasmid (62), a derivative of pSEVA2513 with the constitutive P_EM7_ promoter and the oriV (RSF1010) origin, was replaced with MagFRET from pCMVMagFRET-1 (35) via Gibson assembly (NEBuilder® HiFi DNA Assembly). The primers and plasmids used and the endpoints of the cloned fragments are indicated in Tables S3 and S2, respectively.

### β-galactosidase assay

Overnight cultures grown in N-minimal medium (pH 7.4) supplemented with 10 mM MgCl₂ were diluted into the same medium and grown for 4 hours. The cells were then washed three times with 0.85% NaCl and diluted to an OD₆₀₀ of 0.025 in 25 mL of N-minimal medium (pH 5.5, pH 7.4, or pH 8.5) with the indicated amount of MgCl₂ in a 125 mL baffled flask, followed by incubation at 37°C. After 24 hours, 1 mL of culture was harvested, the cells were pelleted, resuspended in Z buffer, and assayed for β-galactosidase activity in a microtiter plate format, as previously described (63). The β-galactosidase activity units are defined as (μmol of ortho-nitrophenol formed per minute) x10^6^ / (OD_600_ x mL of cell suspension) and are presented as the mean ± standard deviation with n = 6.

### Survival assay

Overnight cultures, grown in N-minimal medium (pH 7.4) supplemented with 10 mM MgCl_2_, were washed with 0.85% NaCl three times, diluted to an OD_600_ value of 0.0375 into 5 mL of N-minimal medium with the indicated amount of MgCl_2_, and incubated at 37°C on a roller drum in a 20 x 150 mm tube. Before and after incubation, serial dilutions of the cultures were plated on LB agar plates that were incubated overnight at 37°C to determine CFU. For the growth curves, the OD_600_ of 250 μL of culture was measured in a BioTek ELx808 Absorbance Reader at the indicated time points. When necessary, the cultures were diluted in the original growth medium to accurately measure the OD_600_ value.

### Mn^2+^ and Co^2+^ sensitivity assay

Overnight cultures, grown in modified MOPS medium (pH 7.4) supplemented with 15 mM glycerol, 1.32 mM K_2_HPO_4_, and 10 mM MgCl₂ were washed three times with 0.85% NaCl, diluted to an OD_600_ value of 0.05 into 10 mL of modified MOPS medium (pH 7.4) supplemented with 15 mM glycerol and with or without 1 mM MnCl_2_ or 0.2 mM CoCl_2_ in 125 mL baffled flask, and incubated at 37°C for 9 hours. Before and after incubation, serial dilutions of the cultures were plated on LB agar plates that were incubated overnight at 37°C to determine CFU.

### Polyamine sensitivity assay

Polyamine stocks were prepared at 500 mM in 0.85% NaCl with HEPES buffer (pH 8.5). Overnight cultures, grown in N-minimal medium (pH 7.4) supplemented with 10 mM MgCl₂ were washed three times with 0.85% NaCl, then diluted 1:200 into 5 mL of N-minimal medium containing either 50 µM or 10 mM MgCl₂. These cultures were incubated at 37°C on a roller drum for 24 hours. Then, cells were washed three times, resuspended in the same volume of 0.85% NaCl with HEPES buffer (pH 8.5), and aliquoted into 1 mL portions in 13 x 100 mm test tubes. The indicated amounts of cadaverine, putrescine, or spermidine were added, and the cultures were incubated at 37°C on a roller drum. At 0 and 24 hours, serial dilutions of the cultures were plated on LB agar plates and incubated overnight at 37°C to determine the CFU.

### Egg white sensitivity assay

Eggs were purchased from the Meat & Egg Sales Room in the Department of Animal Sciences in the University of Illinois at Urbana-Champaign and stored at 4 °C until used (< one month). Egg white was collected from the eggs and mixed with a 4:1 volume of 0.85% NaCl. The mixture was homogenized on ice using a mortar and pestle, and debris was removed by centrifugation. The resulting solution was used as an 80% egg white solution. Note that these egg white solutions did not contain microbes capable of growing on an LB plate. Overnight *Salmonella* cultures grown in N-minimal medium (pH 7.4) supplemented with 10 mM MgCl₂ were diluted into the same medium and grown for 4 hours. The cells were then washed three times with 0.85% NaCl and diluted to an OD₆₀₀ of 0.02 in 1mL of 80% egg white solution or 0.85% NaCl in 13 x 100 mm test tubes. The cultures were incubated at 37°C on a roller drum. At 0 and 24 hours, serial dilutions of the cultures were plated on LB agar plates that were incubated overnight at 37°C to determine the CFU.

### Animal assay

All of the animal work was reviewed and approved by the University of Illinois IACUC and was performed under the protocol number 21197. The competition experiments were performed using 5 to 6-week-old mice. The BALB/cAnNHsd and C3H/HeNHsd mice were purchased from Envigo. The *Salmonella* strains were grown overnight in LB medium, mixed 1:1, and diluted to a target inoculum of approximately 1,000 CFU in 200 μL sterile PBS. The mice were infected via the intraperitoneal route. Each inoculum was plated on LB medium to measure the total inoculum and was replica plated to the appropriate selective medium in order to calculate the input ratio for each strain. After 4 days of infection for the BALB/cAnNHsd mice or 5 days of infection for the C3H/HeNHsd mice, the animals were sacrificed via CO_2_ asphyxiation and cervical dislocation, and their spleens and livers were removed and homogenized. Serial dilutions of the spleen and livers homogenates were plated on LB medium and incubated overnight. The resulting colonies were replica plated to the appropriate selective medium in order to calculate the output ratio for each competition. The competitive index (CI) was calculated as (percent strain A recovered/percent strain B recovered)/(percent strain A inoculated/percent strain B inoculated). The statistical comparisons of individual competitions were done using a Student’s t tests.

### High pH growth assay

Overnight cultures grown in N-minimal medium (pH 7.4), supplemented with 15 mM glycerol and 1 mM MgCl₂, were diluted 800-fold into the same medium and grown for 4.5 hours. The cells were then washed three times with 0.85% NaCl, diluted to an OD₆₀₀ of 0.005 in 10 mL of N-minimal medium (pH 7.4 or 8.5) containing 15 mM glycerol and 1, 10, or 100 mM MgCl₂, and incubated at 37°C in a 125 mL baffled flask. The OD₆₀₀ of culture was measured using a BioTek ELx808 Absorbance Reader at the indicated time points. When necessary, cultures were diluted in the original growth medium to ensure accurate OD₆₀₀ measurements. To determine CFU, serial dilutions of the cultures were plated on LB agar plates and incubated overnight at 37°C.

### Mg^2+^ measurements via inductively coupled plasma-mass spectrometry (ICP-MS)

The cells were cultured as described above for the survival assays. After 7.5 h, cells were collected by centrifugation (7,000 x *g*) and washed twice with 0.85% NaCl. The cell pellets were dried at room temperature and were sent to the University of Georgia Center for Applied Isotope Studies for the determination of the total Mg content. The concentration is reported as μg per g of dry weight and is converted to an intracellular concentration assuming 2.8 x 10^-13^ g per cell (64) and 2.3 x 10^-15^ L per cell (65).

### Free-Mg^2+^ measurements via MagFRET

To obtain a calibration curve, wild-type cells harboring pS2513-MagFRET, pre-grown to mid-exponential phase in N-minimal medium (pH 7.4) with 1 mM MgCl₂, were washed and suspended to an OD₆₀₀ of 0.1 in N-minimal medium (pH 7.4) containing the indicated amounts of MgCl₂, 50 mM sodium benzoate, and 50 mM methylamine HCl. The cells were then incubated at 30°C for 10 minutes. The fluorescence ratio (Citrine_(420/530)_/Cerulean_(420/477)_) was determined using a BioTek Synergy H1 Hybrid Multi-Mode Reader.

To measure free-Mg^2+^ levels, overnight cultures grown in N-minimal medium (pH 7.4), supplemented with 15 mM glycerol and 1 mM MgCl₂, were diluted 750-fold into the same medium and grown for 4.5 hours. The cells were then washed three times with 0.85% NaCl, diluted to an OD₆₀₀ of 0.05 in 12 mL of N-minimal medium (pH 7.4 or 8.5) containing 15 mM glycerol and the indicated amount of MgCl₂, and incubated at 37°C in a 125 mL baffled flask. After 3, 7.5, and 24 hours, aliquots (1.5 mL) were collected, and the fluorescence ratio (Citrine/Cerulean) was determined at an adjusted OD₆₀₀ of 0.1 using a BioTek Synergy H1 Hybrid Multi-Mode Reader. To determine CFU, serial dilutions of the cultures were spotted on LB Km agar plates and incubated overnight at 37°C.

## ACKNOWLEDGMENTS

We are grateful to Jim Imlay and Cari Vanderpool for valuable discussions and Yekaterina Golubeva for critically reading the manuscript. This study was supported by NIH grant R01 AI163687 to JMS. YI performed the experiments. YI and JMS designed the research and wrote the paper. Both authors edited and approved the final version of the manuscript.

