## Supplemental Figs S1-S10, Tables S1-S3 for "The CorC proteins MgpA (YoaE) and CorC protect from excess-cation stress and are required for egg white tolerance and virulence in *Salmonella*"

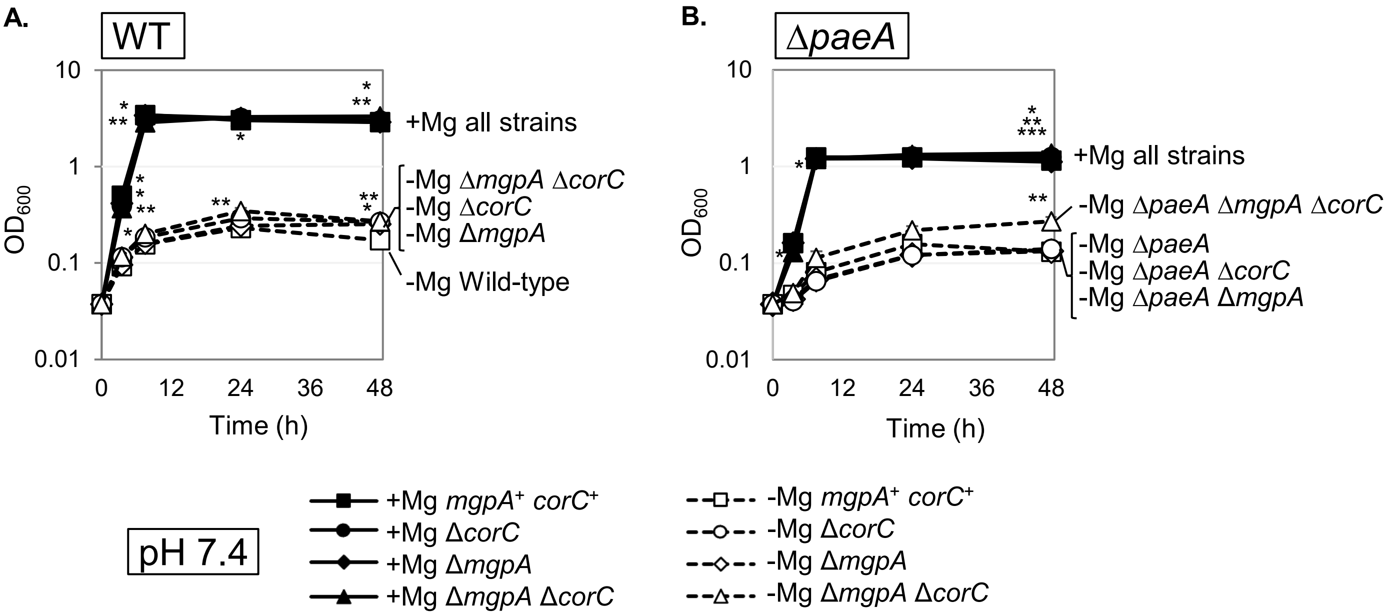

**Fig. S1. $OD_{600}$ of the indicated strains (same experiment as Fig.2) after incubation in $Mg^{2+}$ starvation conditions .** The indicated strains were grown overnight in N-minimal medium pH 7.4 with 10 mM $MgCl_2$, washed, diluted into N-minimal medium pH 7.4 with or without 10 mM $MgCl_2$ (t=0 h), and incubated at 37℃. $OD_{600}$ was determined at the indicated time points in WT (A) and Δ*paeA* (B) backgrounds. Values are mean ± SD, n = 6. Unpaired t test ($p < 0.05$*, 0.005**, 0.0005***) vs *mgpA*$^+$ *corC*$^+$ parent strain at the same timepoint at the same time point and at the same $Mg^{2+}$ concentration. Strains used: 14028, JS2692, JS2693, JS2694, JS2695, JS2696, JS2697, and JS2698.

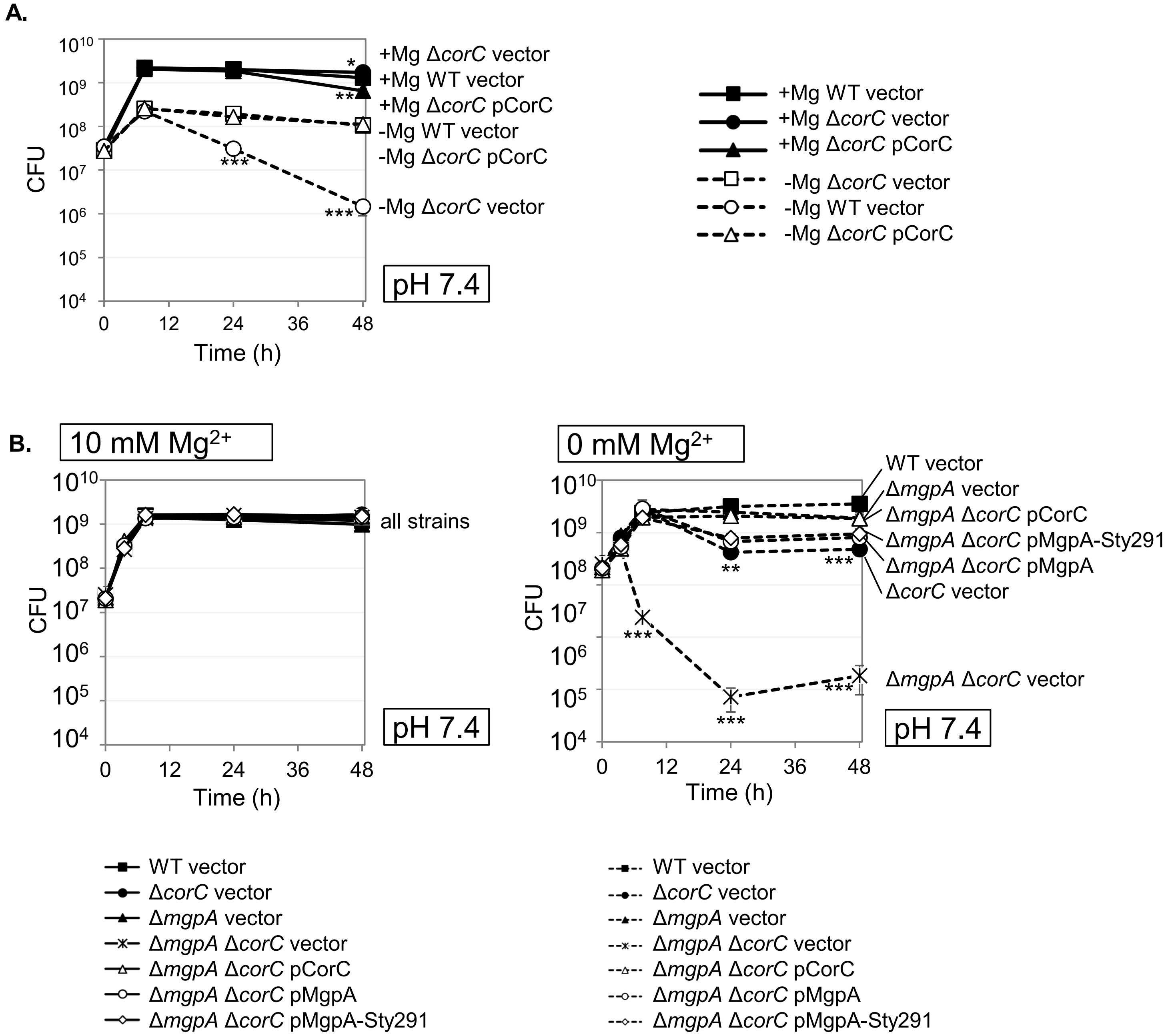

**Fig. S2. Complementation tests in the indicated strains after incubation in $Mg^{2+}$ starvation conditions.** The strains containing pWKS30 plasmid (vector) or pCorC, pMgpA, or pMgpA-Sty291 plasmid as indicated were grown overnight in N-minimal medium pH 7.4 with 10 mM $MgCl_2$, washed, diluted into N-minimal medium pH 7.4 with or without 10 mM $MgCl_2$ (t=0 h), and incubated at 37℃. CFUs were determined at the indicated time points. Values are mean ± SD, n = 6. Unpaired t test ($p$ < 0.05*, 0.005**, 0.0005***) vs WT vector strain at the same time point and at the same $Mg^{2+}$ concentration. Strains used: JS2699, JS2700, JS2701, JS2702, JS2703, JS2704, JS2705, and JS2706

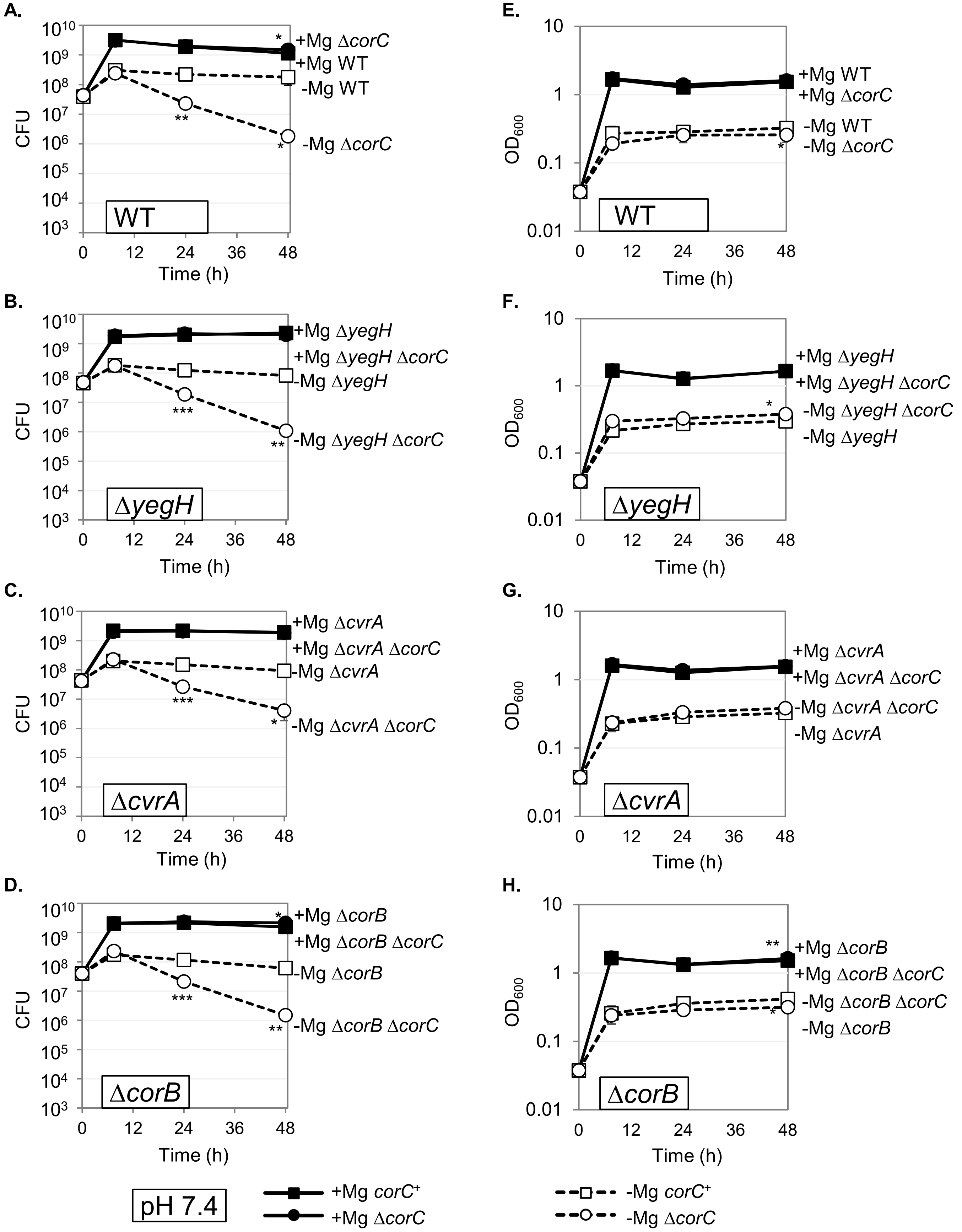

**Fig. S3. YegH, CorB, and CvrA confer no phenotype in stationary phase after $Mg^{2+}$ starvation.** The indicated strains were grown overnight in N-minimal medium pH 7.4 with 10 mM $MgCl_2$, washed, diluted into N-minimal medium pH 7.4 with or without 10 mM $MgCl_2$ (t=0 h), and incubated at 37℃. CFUs and $OD_{600}$ were determined at the indicated time points in WT (A,E), Δ*yegH* (B,F), Δ*cvrA* (C,G), and Δ*corB* (D,H) background. Values are mean ± SD, n = 6. Unpaired t test ($p < 0.05$*, 0.005**, 0.0005***) Δ*corC* vs corresponding *corC*+ parent strain at the same time point and at the same $Mg^{2+}$ concentration. Strains used: 14028, JS2692, JS2707, JS2708, JS2709, JS2710, JS2711, and JS2712.

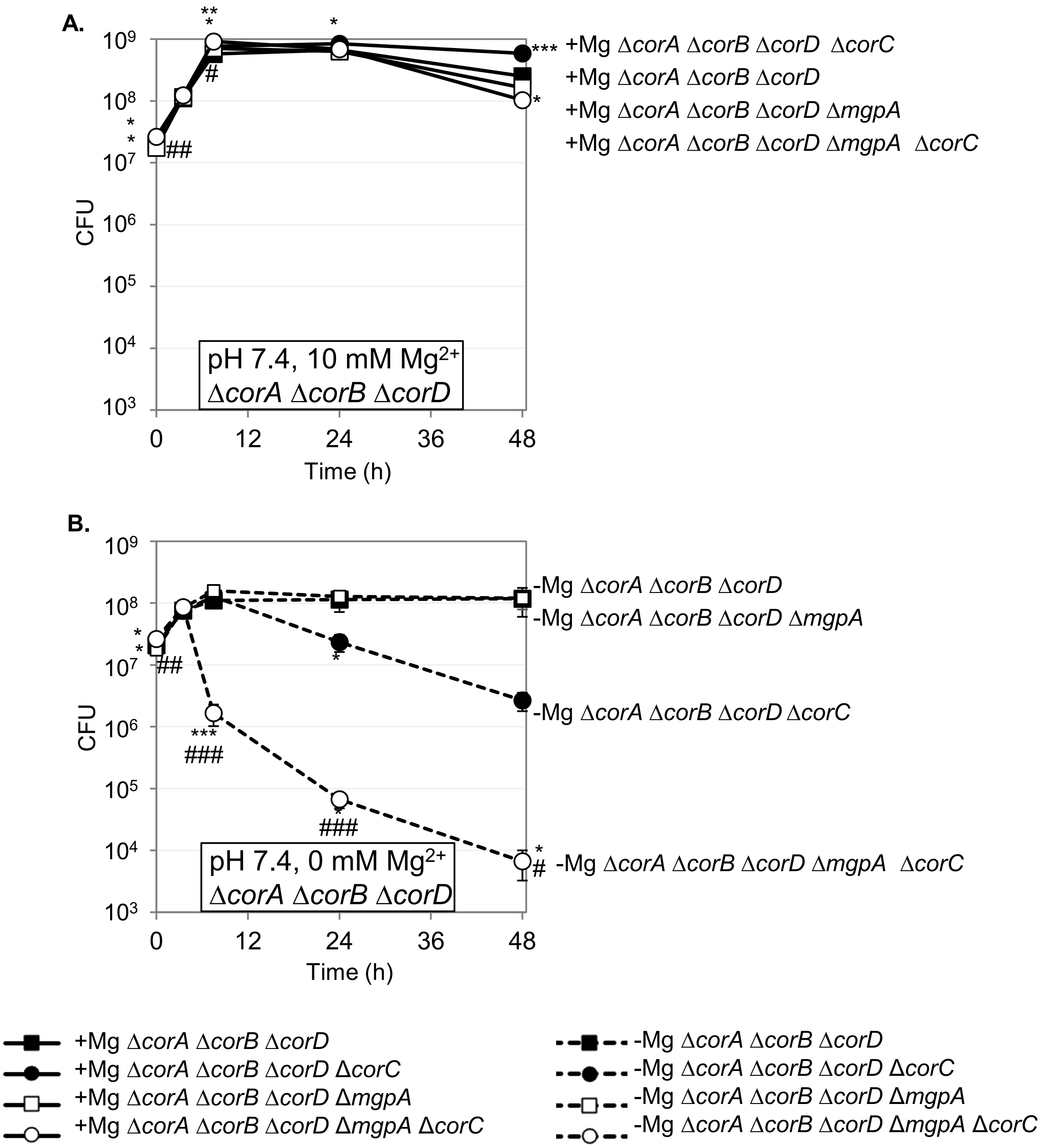

**Fig. S4. Loss of CorA, CorB, and CorD does not affect survival of the ΔcorC and Δ*corC* Δ*mgpA* strains in stationary phase after $Mg^{2+}$ starvation.** The indicated strains were grown overnight in N-minimal medium pH 7.4 with 10 mM $MgCl_2$, washed, diluted into N-minimal medium pH 7.4 (A) with or (B) without 10 mM $MgCl_2$, and incubated at 37℃. CFUs were determined at the indicated time points in the Δ*corA* Δ*corB* Δ*cor*D background. Values are mean ± SD, n = 6. Unpaired t test (p < 0.05*, 0.005**, 0.0005***) versus the control strain at the same time point and at the same $Mg^{2+}$ concentration and (p < $0.05^{\#}$, $0.005^{\#\#}$, $0.0005^{\#\#\#}$) Δ*mgpA corC*+ versus Δ*mgpA* Δ*corC* at the same time point and at the same $Mg^{2+}$ concentration. Strains used: 14028, JS2692, JS2693, JS2694, JS2713, JS2714, JS2715, JS2716, JS2560, JS2717, JS2718, JS2717, JS2720, JS2721, JS2722, and JS2723.

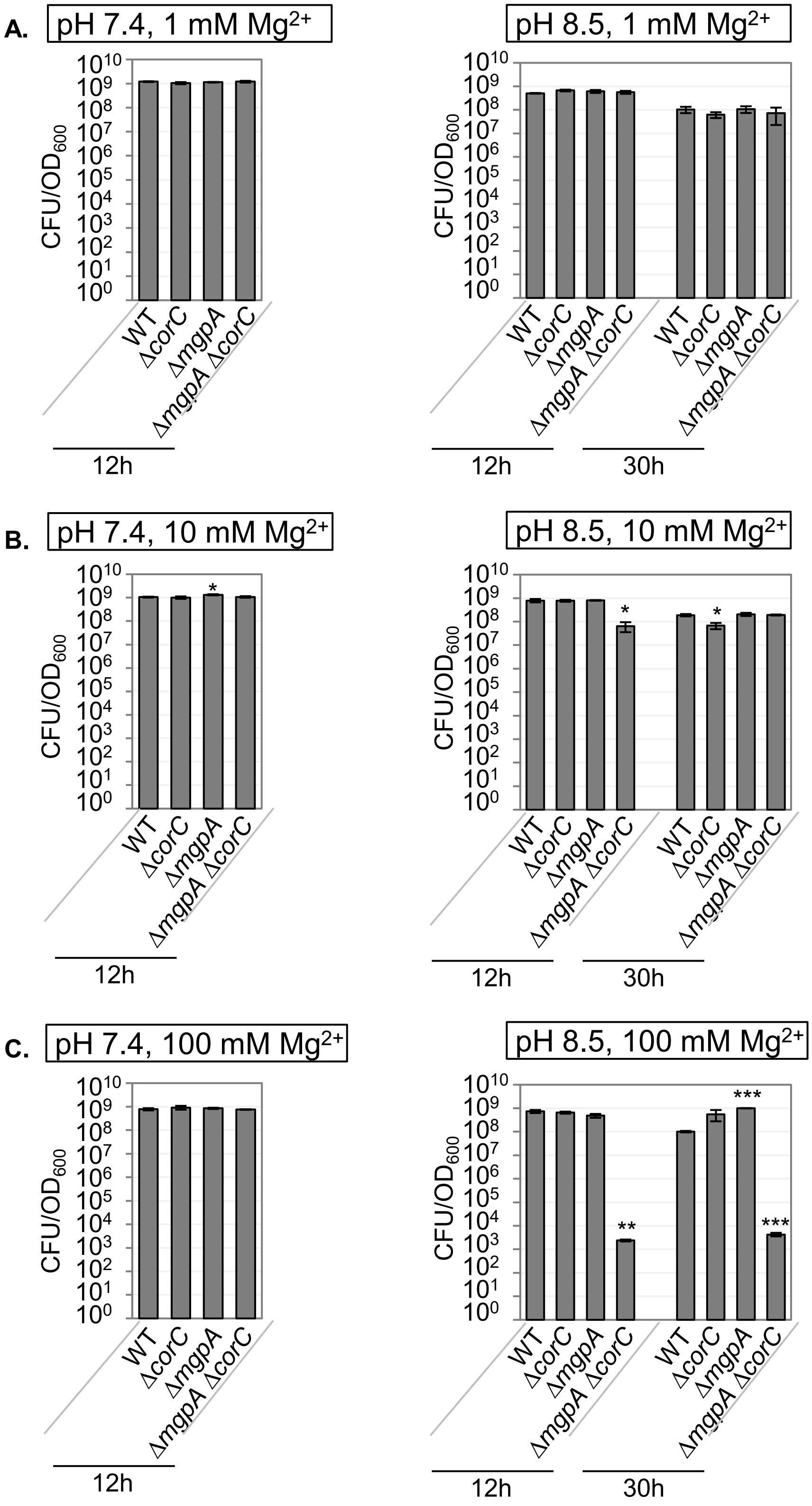

**Fig. S5. The Δ*mgpA* Δ*corC* strain loses viability in the high $Mg^{2+}$ and high pH medium.** The indicated strains were pre-grown to mid-exponential phase in N-minimal medium pH 7.4 with 1 mM $MgCl_2$, washed, and diluted into N-minimal medium pH 7.4 or pH 8.5 with (A) 1 mM, (B) 10 mM, and (C) 100 mM $MgCl_2$ (t=0h), and incubated at 37℃. CFUs were determined at 12h and 30h. Corresponding $OD_{600}$ measurements at the same timepoint from the same experiment are shown in Fig 8. CFUs/$OD_{600}$ were calculated at 12h and 30h. CFU/$OD_{600}$ values are mean ± SD, n = 3. Unpaired t test ($p < 0.05$*, 0.005**, 0.0005***) versus corresponding WT at the same timepoint. Strains used: 14028, JS2692, JS2693, and JS2694.

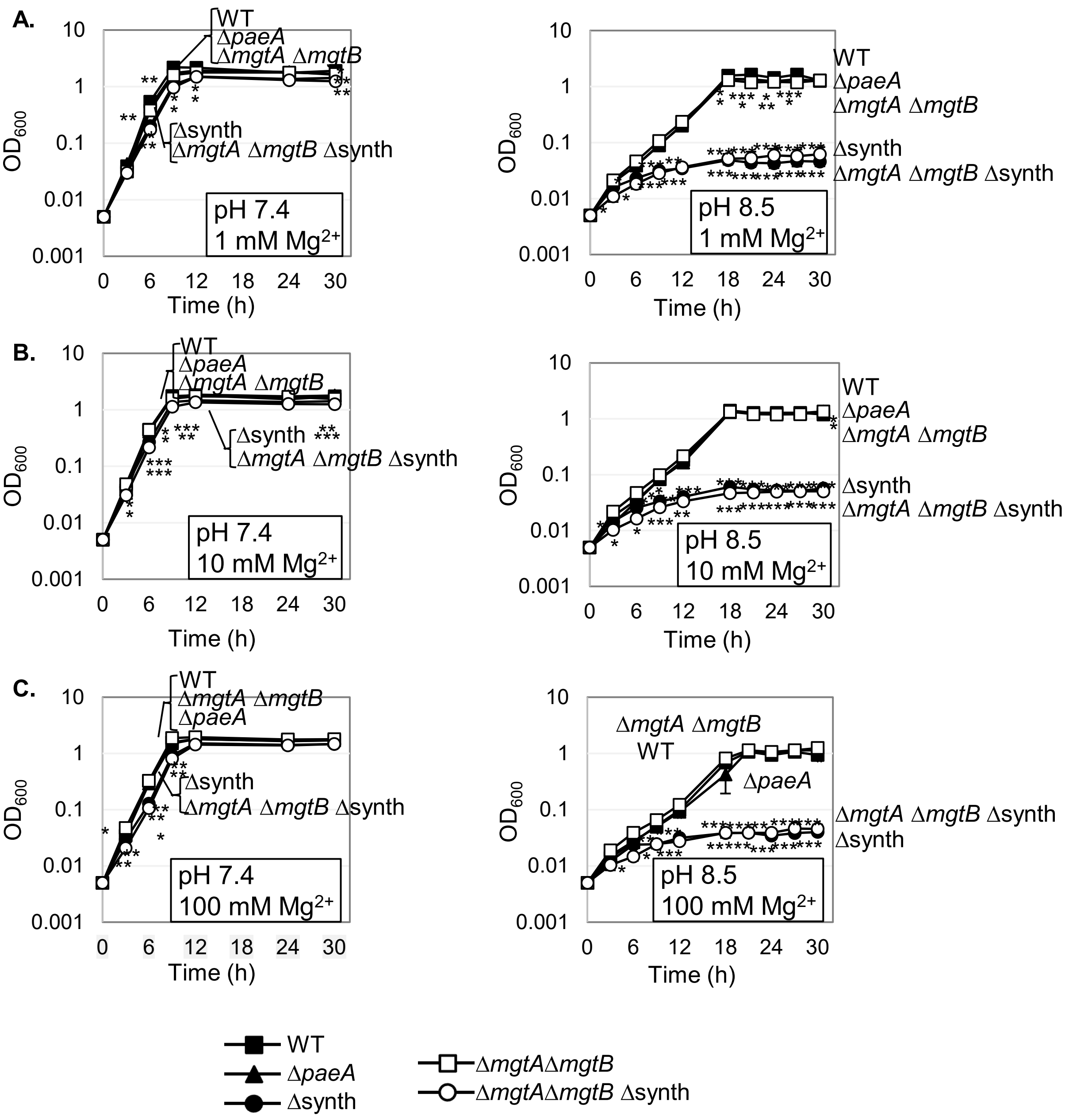

**Fig. S6. pH affects growth under high $Mg^{2+}$ conditions.** The indicated strains were pre-grown to mid-exponential phase in N-minimal medium pH 7.4 with 1 mM $MgCl_2$, washed, and diluted into N-minimal medium pH 7.4 or pH 8.5 with (A) 1 mM, (B) 10 mM, and (C) 100 mM $MgCl_2$ (t=0h), and incubated at 37℃. $OD_{600}$ was determined at the indicated timepoints. Values are mean ± SD, n = 3. Unpaired t test ($p < 0.05$*, 0.005**, 0.0005***) versus corresponding WT at the same timepoint. Strains used: 14028, JS2430, JS2560, JS2562, and JS2563.

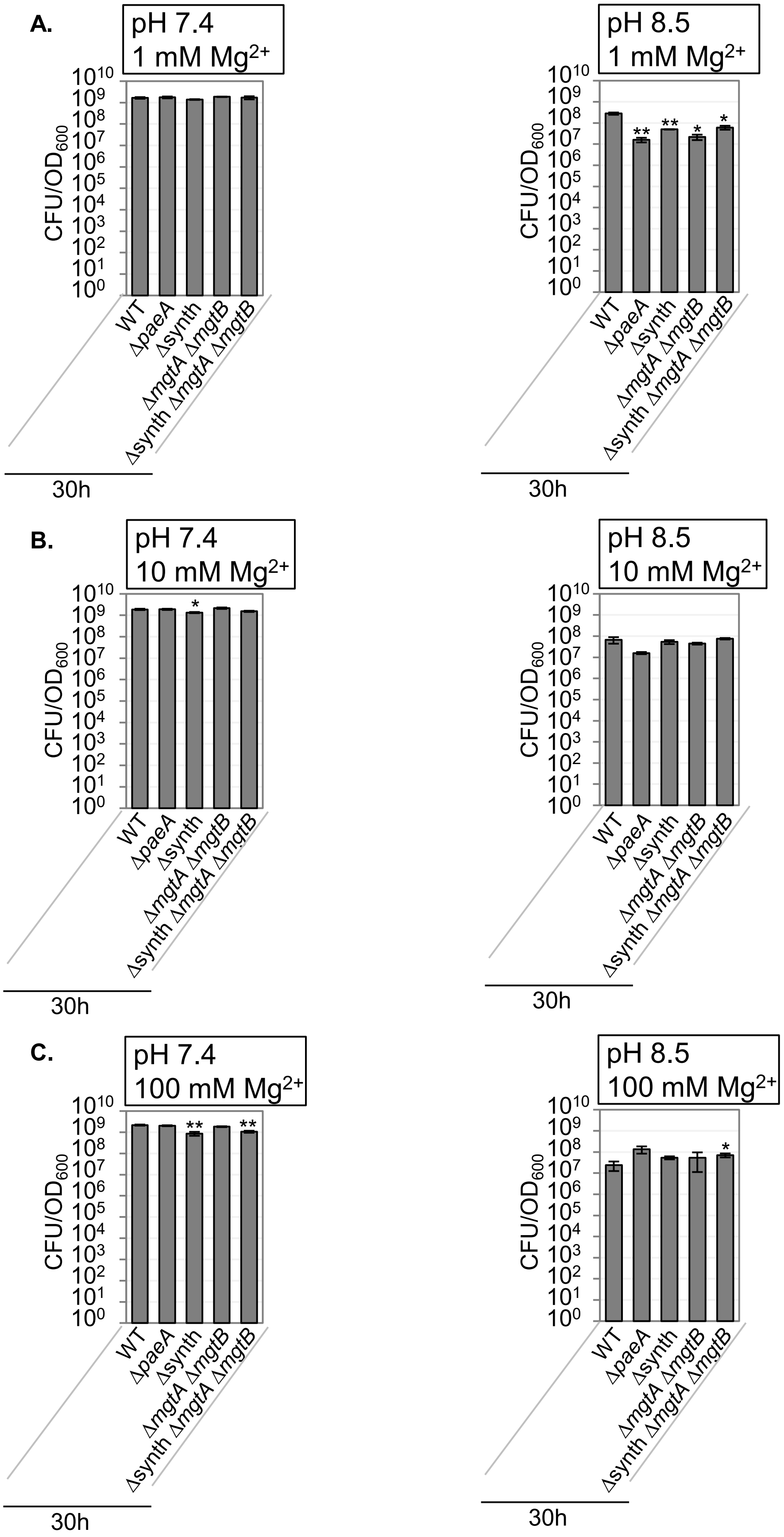

**Fig. S7. The Δ*paeA* strain viability is not affected by the high $Mg^{2+}$ and high pH.** The indicated strains were pre-grown to mid-exponential phase in N-minimal medium pH 7.4 with 1 mM $MgCl_2$, washed, and diluted into N-minimal medium pH 7.4 or pH 8.5 with (A) 1 mM, (B) 10 mM, and (C) 100 mM $MgCl_2$ (t=0h), and incubated at 37℃. CFUs were determined at 30h. Corresponding $OD_{600}$ measurements at the same timepoint from the same experiment are shown in Fig S6. CFUs/$OD_{600}$ were calculated at 30h. CFU/$OD_{600}$ values are mean ± SD, n = 3. Unpaired t test ($p < 0.05$*, 0.005**, 0.0005***) versus corresponding WT at the same timepoint. Strains used: 14028, JS2430, JS2560, JS2562, and JS2563.

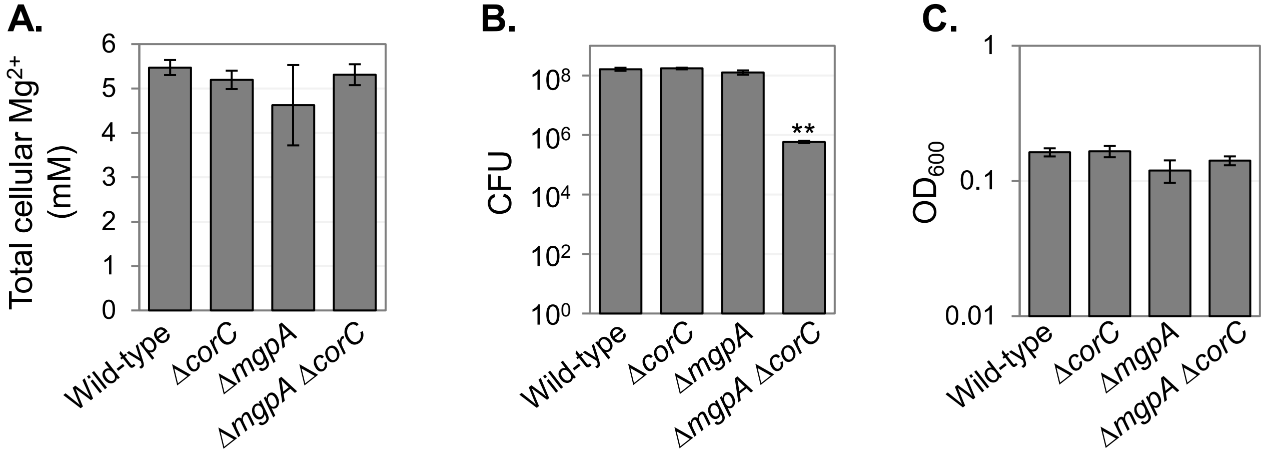

**Fig S8. Total cellular $Mg^{2+}$ levels are not changed by the deletions of *corC* and/or *mgpA* when grown without added Mg.** The indicated strains, overnight cultures grown in N-minimal medium with 10 mM $MgCl_2$, were washed, diluted into N-minimal medium without 10 mM $MgCl_2$, and incubated at 37 ℃ for 7.5 hours. (A) Cellular $Mg^{2+}$ levels were measured via inductively coupled plasma-mass spectrometry (ICP-MS). (B) CFU and (C) $OD_{600}$ were measured. Values are mean ± SD, n = 3. Unpaired t test (*, $P < 0.05$; **, $P < 0.005$; ***, $P < 0.0005$) versus the wild-type. Strains used: 14028, JS2692, JS2693, and JS2694.

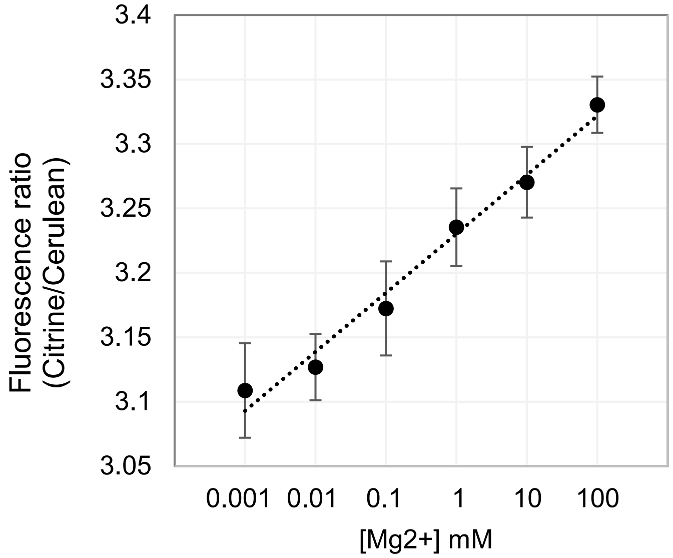

**Fig S9. MagFRET responses to changes in Mg levels in semi-permeabilized *Salmonella* cells.** The wild-type strain harboring pS2513-MagFRET, pre-grown to mid-exponential phase in N-minimal medium (pH 7.4) with 1 mM $MgCl_2$, were washed and diluted to at $OD_{600}$ = 0.1 into N-minimal medium pH 7.4 containing sodium benzoate and methylamine with the indicated amounts of $MgCl_2$, incubated at 30 ℃ for 10 mins, and the fluorescence ratio (Citrine/Cerulean) was determined. Values are mean ± SD, n = 6. Strain used: JS2736.

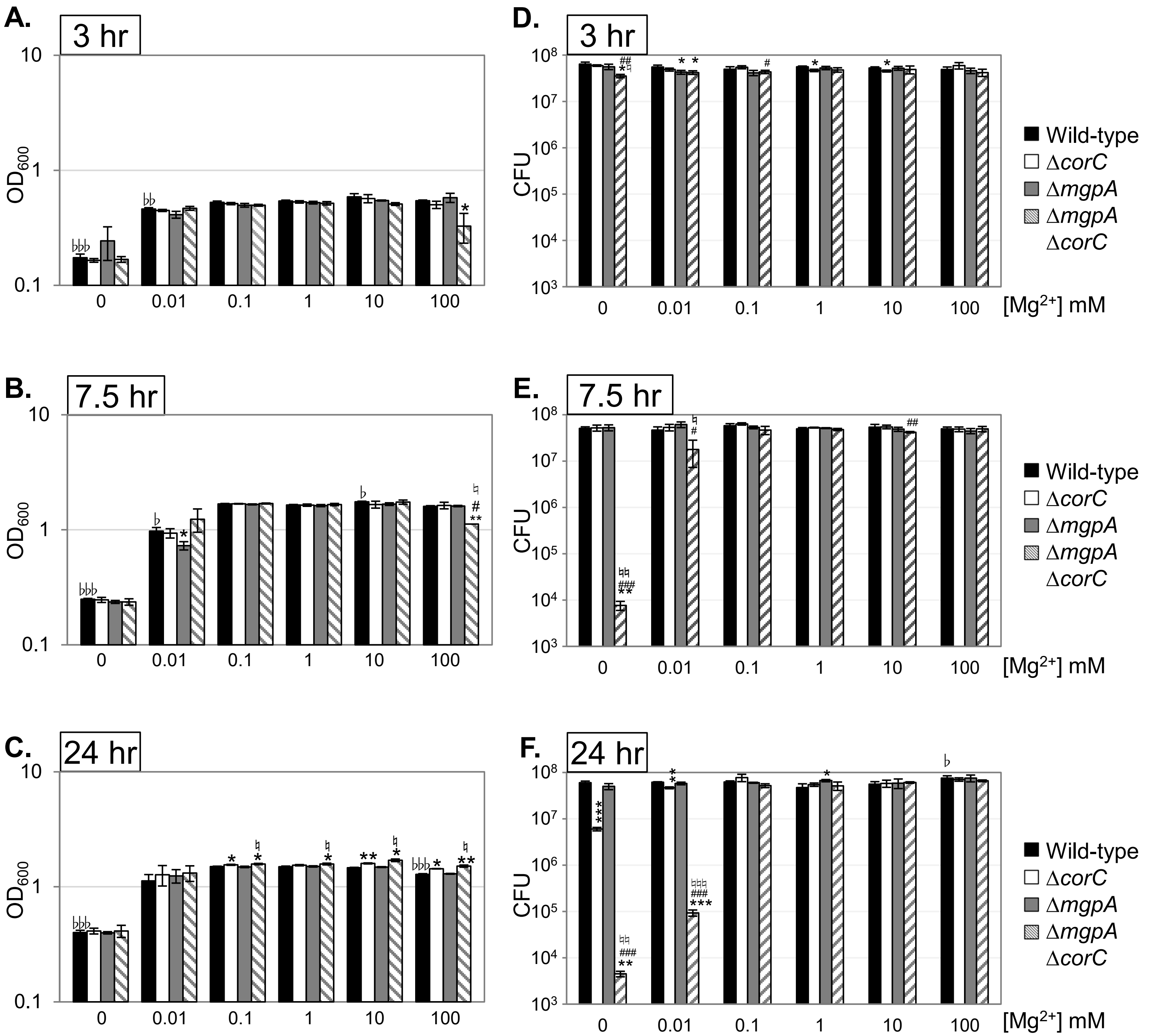

**Fig S10. The Δ*corC* strain shows a lower free-Mg level at stationary phase when grown with 100 mM $MgCl_2$, and the deletion of *mgpA* exacerbates the effect.** The indicated strains harboring pS2513-MagFRET, pre-grown to mid-exponential phase in N-minimal medium (pH 7.4) with 1 mM $MgCl_2$, were washed and diluted into N-minimal medium pH 7.4 with the indicated amounts of $MgCl_2$, and incubated at 37 ℃. Aliquots (1.5 mL) were collected after (A and D) 3, (B and E) 7.5, and (C and F) 24 hours. Corresponding free-Mg levels are shown in Fig 10. (A-C) The $OD_{600}$ of the original culture was shown. (D-F) The CFU of the $OD_{600}$ = 0.1 culture were determined by spotting. Values are mean ± SD, n = 3. Unpaired t test ($P < 0.05$*; $P < 0.005$**; $P < 0.0005$***) versus the wild-type, ($P < 0.05$#; $P < 0.005$##; $P < 0.0005$###) of *mgpA*$^+$ Δ*corC* versus Δ*mgpA* Δ*corC*, or ($P < 0.05$ ♮; $P < 0.005$ ♮♮; $P < 0.0005$ ♮♮♮) of Δ*mgpA* *corC*$^+$ versus Δ*mgpA* Δ*corC* at the same condition. Paired t test ($P < 0.05$ ♭; $P < 0.005$ ♭♭; $P < 0.0005$ ♭♭♭) versus 1 mM $Mg^{2+}$ in the wild-type. Strains used: JS2736, JS2737, JS2738, and JS2739.

Table S1. Bacterial strains

| Strain[a] | Genotype | Endpoint[b] | Ref[c] |
|---|---|---|---|
| 14028 | Wild type | | |
| JS2430 | Δ*paeA*::Cm | | (1) |
| JS2560 | Δ*speED* Δ*speG* Δ*speF* Δ*cadA* Δ*speC* Δ*ldcC* Δ*speB* Δ*speA*::Tc (Δsynth) | | (2) |
| JS2562 | Δ*mgtA*::Kan Δ*mgtB* | | (2) |
| JS2563 | Δ*speED* Δ*speG* Δ*speF* Δ*cadA* Δ*speC* ΔldcC Δ*speB* Δ*speA*::Tc Δ*mgtA*::Kan Δ*mgtB* | | (2) |
| JS2692 | Δ*corC*::Km | 732251-733083 | |
| JS2693 | Δ*mgpA*::Cm | 1937907-1939465 | |
| JS2694 | Δ*mgpA*::Cm Δ*corC*::Km | | |
| JS2695 | Δ*paeA* | | |
| JS2696 | Δ*paeA* Δ*corC*::Km | | |
| JS2697 | Δ*paeA* Δ*mgpA*::Cm | | |
| JS2698 | Δ*paeA* Δ*mgpA*::Cm Δ*corC*::Km | | |
| JS2699 | Wild type + pWKS30 | | |
| JS2700 | Δ*corC*::Km + pWKS30 | | |
| JS2701 | Δ*corC*::Km + pWKS30-*corC* | | |
| JS2702 | Δ*mgpA*::Cm + pWKS30 | | |
| JS2703 | Δ*mgpA*::Cm Δ*corC*::Km + pWKS30 | | |
| JS2704 | Δ*mgpA*::Cm Δ*corC*::Km + pWKS30-*corC* | | |
| JS2705 | Δ*mgpA*::Cm Δ*corC*::Km + pWKS30-*mgpA* | | |
| JS2706 | Δ*mgpA*::Cm Δ*corC*::Km + pWKS30-*mgpA*-sty291 | | |
| JS2707 | Δ*yegH*::Cm | 2254878-2256450 | |
| JS2708 | Δ*yegH*::Cm Δ*corC*::Km | | |
| JS2709 | Δ*cvrA*::Cm | 1910508-1912225 | |
| JS2710 | Δ*cvrA*::Cm Δ*corC*::Km | | |
| JS2711 | Δ*corB*::Cm | | |
| JS2712 | Δ*corB*::Cm Δ*corC*::Km | | |
| JS2713 | Δ*mgtA* Δ*mgtB* | | |
| JS2714 | Δ*mgtA* Δ*mgtB* Δ*corC*::Km | | |
| JS2715 | Δ*mgtA* Δ*mgtB* Δ*mgpA*::Cm | | |
| JS2716 | Δ*mgtA* Δ*mgtB* Δ*mgpA*::Cm Δ*corC*::Km | | |

| | | |
|---|---|---|
| JS2717 | Δ*speED* Δ*speG* Δ*speF* Δ*cadA* Δ*speC* Δ*ldcC* Δ*speB* Δ*speA*::Tc *corC*::Km | |
| JS2718 | Δ*speED* Δ*speG* Δ*speF* Δ*cadA* Δ*speC* Δ*ldcC* Δ*speB* Δ*speA*::Tc *mgpA*::Cm | |
| JS2720 | Δ*corA* | 4171866-4172816 |
| JS2721 | Δ*corA* Δ*corC*::Km | |
| JS2722 | Δ*corA* Δ*mgpA*::Cm | |
| JS2723 | Δ*corA* Δ*mgpA*::Cm Δ*corC*::Km | |
| JS2724 | Δ*paeA* Δ*corC* | |
| JS2725 | Δ*paeA* Δ*mgpA* | |
| JS2726 | Δ*paeA* Δ*corC* Δ*mgpA* | |
| JS2728 | Δ*corC* | |
| JS2729 | Δ*mgpA* | |
| JS2730 | Δ*mgpA* Δ*corC* | |
| JS2731 | Δ*mgpA* Δ*corC* + pWKS30 | |
| JS2732 | Δ*mgpA* Δ*corC* + pWKS30-*corC* | |
| JS2733 | Δ*mgpA* Δ*corC* + pWKS30-*mgpA* | |
| JS2734 | att::*corC*60-lacZ | 733046-734983 |
| JS2735 | att::*mgpA*24-lacZ | 1939442-1940465 |
| JS2736 | 14028 + pS2513-MagFRET1 | |
| JS2737 | Δ*corC* + pS2513-MagFRET1 | |
| JS2738 | Δ*mgpA* + pS2513-MagFRET1 | |
| JS2739 | Δ*mgpA* Δ*corC* + pS2513-MagFRET1 | |

a All *Salmonella* strains are isogenic derivatives of *S. enterica* serovar Typhimurium strain 14028.

b Numbers indicate the base pairs that are deleted (inclusive) as defined in the *S. enterica* serovar Typhimurium 14028 genome sequence (National Center for Biotechnology Information; NC_016856.1)

c This study, unless otherwise indicated

Table 2. Plasmids

| Name | Characteristic | Cloned endpoint[a] | Ref[b] |
|---|---|---|---|
| pKD46 | *bla* PBAD *gam bet exo* pSC101 *oriTS* | | (3) |
| pCP20 | *bla cat* cI857 λPR *flp* pSC101 *oriTS* | | (3) |
| pKD3 | *bla* FRT *cat* FRT PS1 PS2 *oriR6K* | | (3) |
| pKD4 | *bla* FRT *aph* FRT PS1 PS2 *oriR6K* | | (3) |
| pWKS30 | *bla* pSC101 *ori* | | (4) |
| pWKS30-*corC* | | 732218-733301 | |
| pWKS30-*mgpA* | | 1937879-1939936 | |
| pWKS30-*mgpA*-sty291 | | 1937879-1939936 | |
| pDX1 | *lacZ* tL3 λ*attP oriR6K aacIV tmgB* | | (5) |
| pDX1-*corC*-lacZ | | 733046-734983 | |
| pDX1-*mgpA*-lacZ | | 1939442-1940465 | |
| pS2513-PHP | | | (6) |
| pCMVMagFRET-1 | | | (7) |
| pS2513-MagFRET1 | | | |

a Numbers indicate the base pairs that are cloned (inclusive) as defined in the *S. enterica* serovar Typhimurium 14028 genome sequence (National Center for Biotechnology Information; NC_016856.1)

b This study, unless otherwise indicated

Table S3. Primers used

| Name | Sequence (5' to 3’) | Usage |
|---|---|---|
| YIV_pWKS30-12 | ATTGCGTTGCGCTCACTGCC | pWKS30-*corC*, pWKS30-*mgpA*, and pWKS30-*mgpA*-sty291 |
| YIV_pWKS30-13 | AACGTCGTGACTGGGAAAACC | pWKS30-*corC*, pWKS30-*mgpA*, and pWKS30-*mgpA*-sty291 |
| YIS_73-18 | GTTTTCCCAGTCACGACGTTCTATTGCTGTTATTCGTCCAGTTTTG | pWKS30-*corC* |
| YIS_73-29 | GGCAGTGAGCGCAACGCAATATGAGGATCCGTACATTGCC | pWKS30-*corC* |
| YIS_193-7(RE) | GGCAGTGAGCGCAACGCAATTGCTACCTCCTTTATTATTGTCAACAC | pWKS30-*mgpA* and pWKS30-*mgpA*-sty291 |
| YIS_193-5 | GTTTTCCCAGTCACGACGTTCTATAGACTTGATTCCTGCGTG | pWKS30-*mgpA* |
| YIS_193-6 | GTTTTCCCAGTCACGACGTTCCATTAGCGAATTTCCACAGTG | pWKS30-*mgpA*-sty291 |
| YIS_pDX1-1 | ATGACCATGATTACGGATTC | pDX1-*corC*-lacZ and pDX1-*mgpA*-lacZ |
| YIS_pDX1-3 | TTGGATCCTCTAGAGTCGACCTGCAG | pDX1-*corC*-lacZ and pDX1-*mgpA*-lacZ |
| YIS_73-11 | CTGCAGGTCGACTCTAGAGGATCCAACCCTACTGAAAAGGCAATAATG | pDX1-*corC*-lacZ |
| YIS_73-44 | GAATCCGTAATCATGGTCATGGAAAAAAATCCCTTTTTAC | pDX1-*corC*-lacZ |
| YIS_194-9 | CTCTAGAGGATCCAACTGACCGTGAATGAGGCGGTC | pDX1-*mgpA*N24-lacZ |
| YIS_193-26 | CGTAATCATGGTCATTGAGGGATCCATTAATAATTCCATGAC | pDX1-*mgpA*N24-lacZ |
| YIV_pS2513N | ATGTTTTTCCTCCTAAGCTT | pS2513-MagFRET1 |
| YIV_pS2513C | ACTAGTCTTGGACTCCTGTT | pS2513-MagFRET1 |
| YIV_MagFRET1N | TAGGAGGAAAAACATATGGGCCATATGGTG | pS2513-MagFRET1 |
| YIV_MagFRET1C | GAGTCCAAGACTAGTTTACTTGTACAGCTCGTC | pS2513-MagFRET1 |
